# Mitochondrial-Encoded Peptide MOTS-c is an Exercise-Induced Regulator of Aging Metabolic Homeostasis and Physical Capacity

**DOI:** 10.1101/2019.12.22.886432

**Authors:** Joseph C. Reynolds, Rochelle W. Lai, Jonathan S.T. Woodhead, James H. Joly, Cameron J. Mitchell, David Cameron-Smith, Ryan Lu, Pinchas Cohen, Nicholas A. Graham, Bérénice A. Benayoun, Troy L. Merry, Changhan Lee

## Abstract

Healthy aging can be promoted by enhancing metabolic fitness and physical capacity (*1, 2*). Mitochondria are chief metabolic organelles with strong implications in aging (*3–8*). In addition to their prominent role in bioenergetics, mitochondria also coordinate broad physiological functions by communicating to other cellular compartments or distal cells using multiple factors (*9, 10*), including peptides that are encoded within their own independent genome (*11, 12*). However, it is unknown if aging is actively regulated by factors encoded in the mitochondrial genome. MOTS-c is a mitochondrial-encoded peptide that regulates metabolic homeostasis (*13, 14*), in part, by translocating to the nucleus to regulate adaptive nuclear gene expression in response to cellular stress (*15–17*). Here, we report that MOTS-c is an exercise-induced mitochondrial-encoded peptide that significantly enhanced physical performance when administered to young (2 mo.), middle-aged (12 mo.), and old (22 mo.) mice. In humans, we found that endogenous MOTS-c levels significantly increased in response to exercise in skeletal muscle (11.9-fold) and in circulation (1.5-fold). Systemic MOTS-c treatment in mice significantly enhanced the performance on a treadmill of all age groups (~2-fold). MOTS-c regulated (i) nuclear genes, including those related to metabolism and protein homeostasis, (ii) glucose and amino acid metabolism in skeletal muscle, and (iii) myoblast adaptation to metabolic stress. Late-life (23.5 mo.) initiated intermittent MOTS-c treatment (3x/week) improved physical capacity and trended towards increasing lifespan. Our data indicate that aging is regulated by genes that are encoded not only in the nuclear genome (*18, 19*), but also in the mitochondrial genome. Considering that aging is the major risk factor for multiple chronic diseases (*20, 21*), our study provides new grounds for further investigation into mitochondrial-encoded regulators of healthy lifespan.

## Main Text

Organismal fitness requires continuous adaptive cellular stress responses to the ever-shifting internal and external environment. The capacity to maintain metabolic homeostasis declines with age, which impedes parenchymal function and ultimately diminishes physical capacity. Mitochondria not only produce the bulk of cellular energy, but also coordinate adaptive cellular homeostasis by dynamically communicating to the nucleus (*9*) and other subcellular compartments (*22*). Mitochondrial communication is mediated by multiple nuclear-encoded proteins, transient molecules, and mitochondrial metabolites (*23*).

Mitochondria possess a distinct circular genome that has been traditionally known to host only 13 protein-coding genes. However, short open reading frames (sORFs) encoded in the mitochondrial genome have been recently identified. Such sORFs produce bioactive peptides, collectively referred to as mitochondrial-derived peptides (MDPs), with broad physiological functions (*11, 12*). MOTS-c (mitochondrial ORF of the 12S rDNA type-c) is an MDP that promotes metabolic homeostasis, in part, via AMPK (*13, 14*) and by directly regulating adaptive nuclear gene expression following nuclear translocation (*15, 16*). MOTS-c expression is age-dependent and detected in multiple tissues, including skeletal muscle, and in circulation (*13, 14, 24*), thus it has been dubbed a “mitochondrial hormone” (*14*) or “mitokine” (*25, 26*). In fact, systemic MOTS-c treatment reversed diet-induced obesity and diet- and age-dependent insulin resistance in mice (*13*). We tested if MOTS-c functions as a mitochondrial-encoded regulator of physical capacity and performance (*2, 27, 28*) in young (2 mo.), middle-aged (12 mo.), and old (22 mo.) mice.

To determine if endogenous MOTS-c responds to physical exertion, and thus may be involved in driving adaptation to enhance physical capacity, we collected skeletal muscle and plasma from sedentary healthy young male volunteers (24.5 ± 3.7 years old and BMI 24.1 ± 2.1) that exercised on a stationary bicycle (Fig. 1A). Samples were collected before, during (plasma only), and after exercise and following a 4-hour rest. Western blotting revealed that endogenous MOTS-c levels in skeletal muscle significantly increased after exercise (11.9-fold) and remained elevated after a 4-hour rest (18.9-fold) (Fig. 1B, C). ELISA revealed that circulating endogenous MOTS-c levels also significantly increased during (1.6-fold) and after (1.5-fold) exercise, which then returned to baseline after 4 hours of resting (Fig. 1D, fig. S1). These findings suggest that exercise induces the expression of mitochondrial-encoded regulatory peptides in humans.

**Fig. 1.**
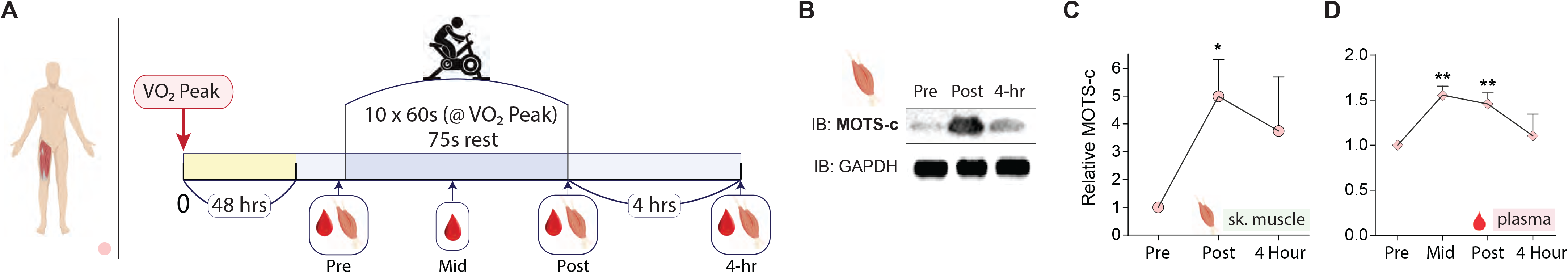
MOTS-c responds to and regulates exercise in young subjects. (**A**) Schedule of exercise on a stationary bicycle and blood and skeletal muscle collection in young male subjects (n=10). (**B, C**) Representative western blot of MOTS-c from skeletal muscle and quantification (**D**) Quantification of serum MOTS-c levels by ELISA. Data expressed as mean +/− SEM. Wilcoxon matched-pairs signed rank test was used for (**C, D**).

We next probed if MOTS-c functions as an exercise-induced mitochondrial signal that improves physical capacity by treating young mice (CD-1; outbred) daily with MOTS-c [5mg/kg/day; intraperitoneal injections (IP)] for 2 weeks. The rotarod performance test, whereby mice are placed on a rotating rod, revealed that daily MOTS-c significantly improved physical capacity (fig. S2A), but not grip strength (fig. S2B) in young mice. Because the rotarod test can also be affected by cognitive capacity, we assessed learning and memory using the Barnes maze and found no improvement (fig. S2C, D).

A treadmill running test confirmed that MOTS-c treatment can enhance physical performance. Because MOTS-c is regulator of metabolic homeostasis that prevented high-fat diet (HFD)-induced obesity and insulin resistance (*13*), we tested if MOTS-c also improved running performance under metabolic (dietary) stress. We fed young mice (CD-1) a HFD (60% calories from fat) and treated them with 2 doses of MOTS-c (5 and 15 mg/kg/day; IP) (fig. S3A). Mice on the higher dose of MOTS-c showed significantly superior running capacity following 10 days of treatment (Fig. 2A-C), but not 7 days of treatment (fig. S4A). We progressively increased the treadmill speed to test both endurance and speed. The final stage, which required mice to sprint (23m/s), was reached by 100% of mice on the higher dose of MOTS-c, but only 16.6% in the lower dose and control (vehicle) groups (Fig. 2D). Body composition analysis using a time-domain NMR analyzer revealed that both doses of MOTS-c significantly retarded fat gain and that the high dose significantly increased lean mass in young mice (CD-1) (fig. S5A-C), in accord with prior reports (*13*).

**Fig. 2.**
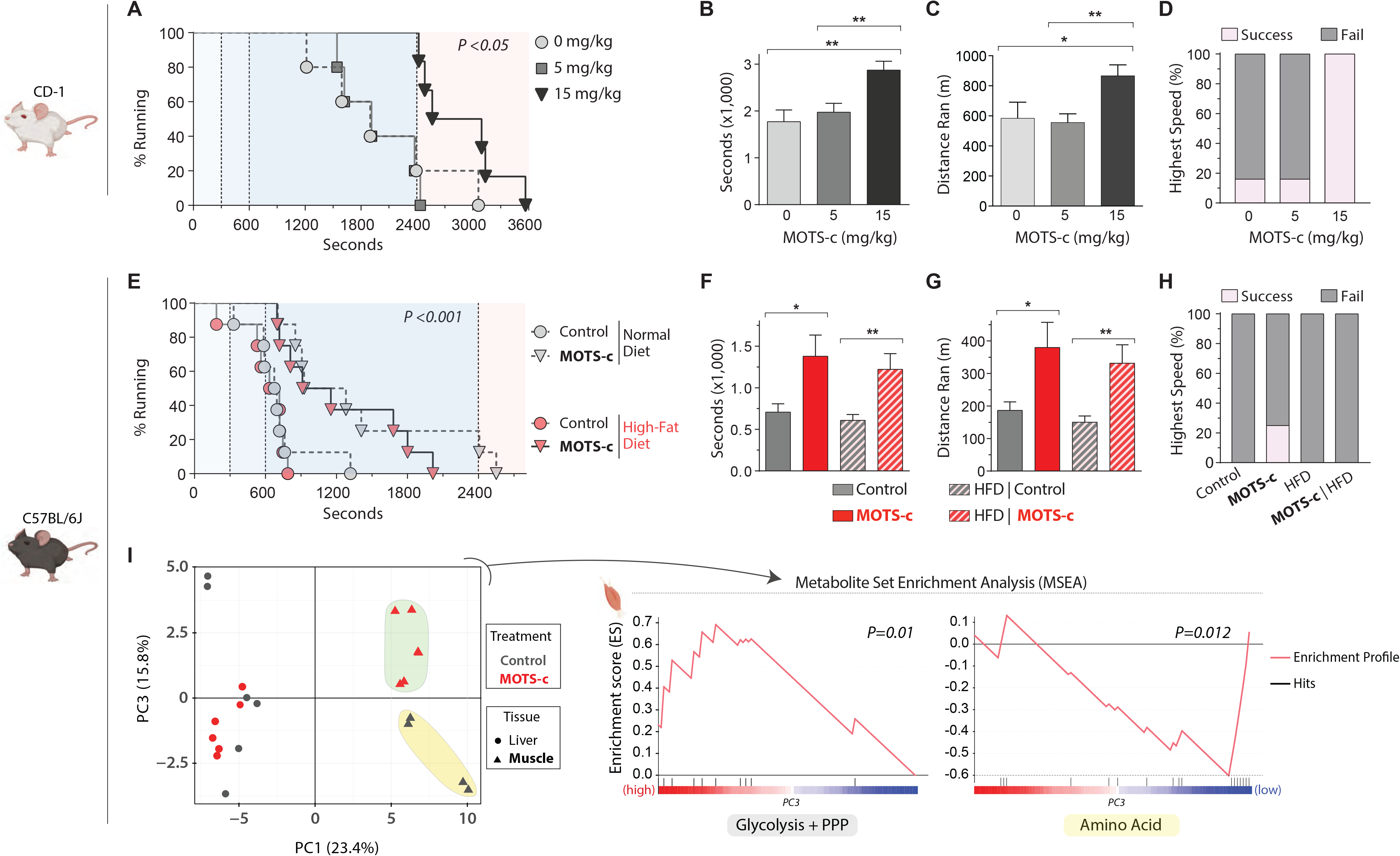
MOTS-c treatment increases physical capacity in young mice regardless of diet. (**A-C**) Treadmill performance of 12-week old male CD-1 (outbred) mice fed a normal diet (n=5-6); (**A**) running curves, (**B**) total time on treadmill, (**C**) total distance ran, and (**D**) percent capable of reaching the highest speed (sprint). (**E-H**) Treadmill performance of 12-week old male C57BL/6J (inbred) mice fed a HFD (n=8); (**E**) running curves, (**F**) total time on treadmill (**G**) total distance ran (**H**) percent capable of reaching the highest speed (final stage). (**I**) PCA and MSEA on metabolomic data from skeletal muscle and liver of C57BL/6J mice that were fed a HFD, treated with MOTS-c, and exercised. Data expressed as mean +/− SEM. Log-rank (Mantel-Cox) test was used for (**A, E**). Otherwise, all statistics were performed using the Student’s *t*-test. \**P*<0.05, ** *P*<0.01, *** *P*<0.001.

In young CD-1 mice, we simultaneously initiated MOTS-c treatment and a HFD (fig. S3A). To test if MOTS-c can improve physical performance in mice that have been on a HFD, we fed young C57BL/6J mice a HFD, or a normal diet, for 2 weeks before initiating daily MOTS-c injections (15 mg/kg/day) for 2 weeks prior to a treadmill running test (fig. S3B). MOTS-c treatment significantly enhanced running performance on the treadmill regardless of the diet (Fig. 2E-G, fig. S4B). MOTS-c treatment enabled 25% of the young C57BL/6J mice to enter the final running stage (highest speed) on a normal diet, but none on a HFD (Fig. 2H). Consistent with our prior study (*13*), MOTS-c treatment curbed HFD-induced weight gain in C57BL/6J mice (fig. S5D), which was largely driven by reduced fat accumulation (fig. S5E), but not loss of lean mass (fig. S5F), as determined by an NMR-based body composition analysis. Further, targeted metabolomics revealed that MOTS-c treatment significantly regulated (i) glycolysis/PPP (pentose phosphate pathway) and (ii) amino acid metabolism (Fig. 2I, fig. S3B) in skeletal muscle, but not in liver, consistent with our previous study (*13*). Together, these data indicate that MOTS-c treatment can improve overall physical performance, in part, by targeting skeletal muscle metabolism in young mice.

Aging is accompanied by a progressive decline in mitochondrial function (*1, 8*) and loss of metabolic homeostasis, in which MOTS-c may play a role (*2, 9*). Aging is associated with reduced MOTS-c levels in certain tissues, including the skeletal muscle, and in circulation (*13, 24*). We previously showed that an acute one-week MOTS-c treatment reversed age-dependent insulin resistance in mouse skeletal muscle (*13*). Thus, we investigated if promoting metabolic homeostasis by MOTS-c treatment could reverse age-dependent decline in physical capacity. Middle-aged (12 mo.) and old (22 mo.) C57BL/6N mice were treated daily with MOTS-c (15 mg/kg/day; IP) for 2 weeks, then subjected to a treadmill running test (Fig. 3A). Both middle-aged and old mice ran significantly longer following MOTS-c treatment (Fig. 3B). Old mice ran longer (2-fold) (Fig. 3C) and farther (2.16-fold) (Fig. 3D) when treated with MOTS-c. Further, MOTS-c enabled 17% of the old mice to enter the final running stage (highest speed), whereas none in the untreated group were successful (Fig. 3E). Notably, MOTS-c treatment enabled old mice to outperform untreated middle-aged mice, suggesting a more pervasive physical re-programming rather than just rejuvenation. Respiratory exchange ratio (RER), measured using a metabolic cage, indicates fuel preference (1.0: carbohydrates, 0.7: fat). “Metabolic flexibility”, which refers to the overall adaptive capacity to a shift in metabolic supply-demand equilibrium (*e.g.* exercise), declines with age (*29, 30*). Indeed, old mice relied on carbohydrates regardless of the time of day (Fig. 3F), whereas middle-aged mice, and MOTS-c-treated old mice, exhibited a circadian-dependent shift in fuel usage that favored fat during the daytime (Fig. 3F), coinciding with the low-feeding hours (fig. S6). Metabolomic analysis on skeletal muscle collected immediately post-exercise (a 30-minute run at a fixed moderate speed) in MOTS-c-treated (2 weeks) mice revealed that MOTS-c significantly regulated glycolysis and amino acid metabolism (Fig. 3G); the skeletal muscles of non-exercised mice did not show significant alterations in response to MOTS-c (fig. S7), suggesting that MOTS-c induces an adaptive metabolic response to exercise. To begin to understand the molecular mechanisms underlying the effects of MOTS-c, we performed RNA-seq analysis on the same skeletal muscles used for metabolomics. Although individual-to-individual variability was high, Gene Set Enrichment Analysis (GSEA) using the KEGG pathway database revealed that MOTS-c regulated processes related to (i) metabolism, including those known to be regulated by MOTS-c (*e.g.* AMPK signaling, glycolysis, and central carbon metabolism) (*13, 15*), and (ii) longevity (FDR < 15%; select pathways in Fig. 3H; full analysis in able S1). Gene Ontology Biological Process (GO_BP) analysis revealed a broader range of processes, including metabolism (lipid, carbohydrate, amino acid, and nucleotides), oxidative stress response, immune response, and nuclear transport (FDR < 15%; select pathways in fig. S8; full analysis in table S2), again, consistent with our previous studies (*13, 15*). The rotarod performance test confirmed that MOTS-c treatment improved physical capacity in old mice (fig. S9A), while learning and memory was not affected as determined using the Y-maze test (fig. S9B), consistent with our observations in young mice (fig. S2). Together, these data suggest that MOTS-c treatment can significantly improve physical capacity in old mice, in part, by regulating skeletal function and improving “metabolic flexibility”.

**Fig. 3.**
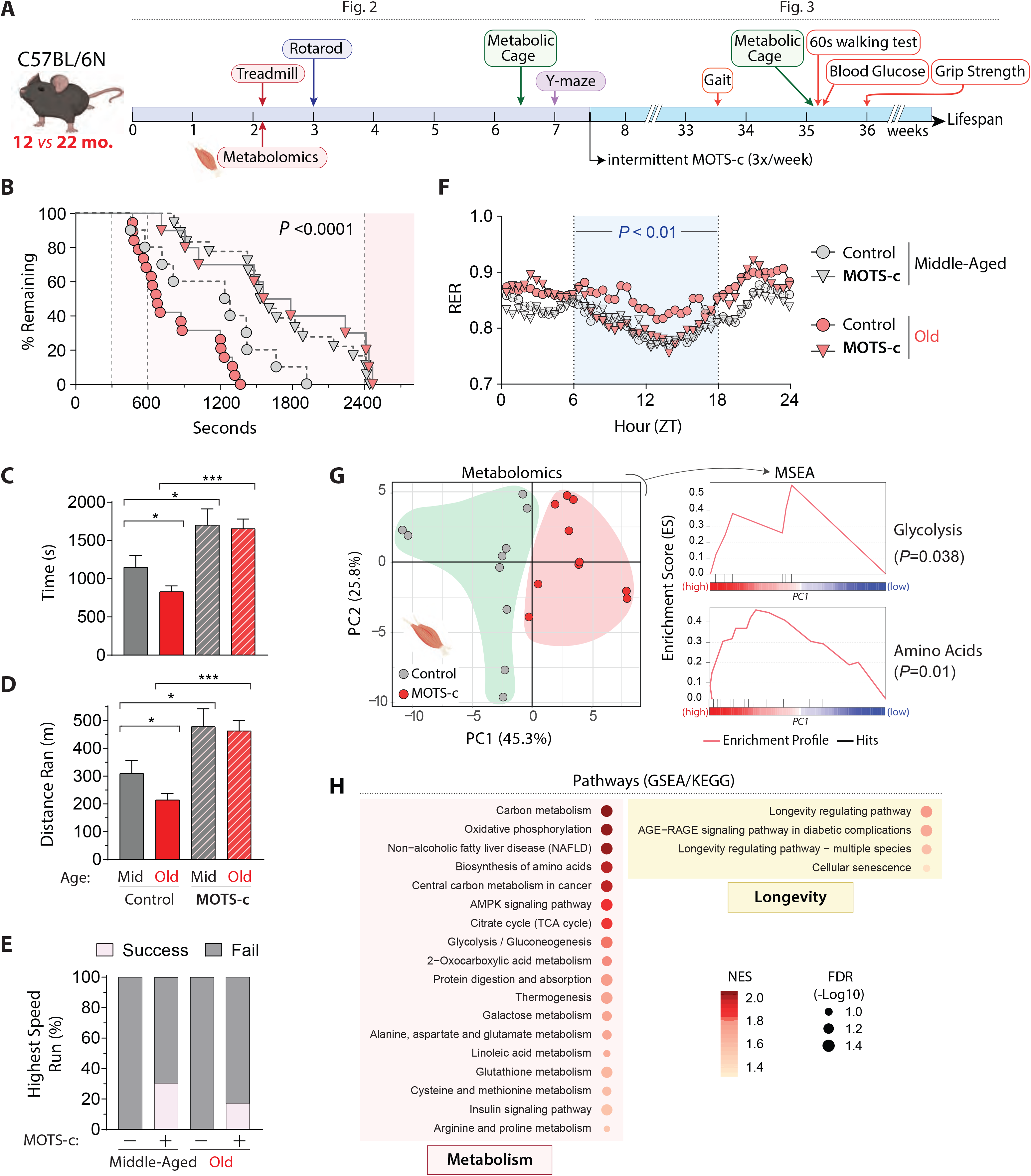
Acute MOTS-c treatment enhances physical capacity in old mice. (**A**) Schedule of MOTS-c treatment and assays in middle-aged and old C57BL/6N mice (n=10 and 16-19, respectively), including (**B**) treadmill running curves, (**C**) total time on treadmill, (**D**) total distance ran on treadmill, and (**E**) percent capable of reaching the highest speed on a treadmill (final stage). (**F**) Respiratory exchange ratio (RER) following 2 weeks of daily MOTS-c injection (n=4). (**G, H**) Skeletal muscle from treadmill-exercised old mice (22.5 months) treated daily with MOTS-c (15 mg/kg/day) for 2 weeks (n=10) were subject to (**G**) metabolomics and analyzed using PCA and MSEA and (**H**) GSEA analysis of muscle RNA-seq analysis. Balloon plots of select enriched terms using Gene Ontology Biological Process (GO_BP) database at false discovery rate (FDR) < 15%. Full GSEA results are available in table S1. Data expressed as mean +/− SEM. Log-rank (Mantel-Cox) test was used for (**B**) and two-way ANOVA (repeated measures) was used for (**F**). GSEA statistics from R package ‘clusterProfiler’ were used for (**H**). Otherwise, all statistics were performed using the Student’s *t*-test. \**P*<0.05, ** *P*<0.01, *** *P*<0.001.

Age is the major risk factor for many chronic diseases and interventions that delay aging may extend healthy lifespan (*20, 31–33*). Anti-aging interventions that are applied later in life would be more translationally feasible compared to life-long treatments (*34, 35*). Building on the treadmill running tests, we tested if a late-life initiated (~24 mo.) intermittent (LLII) MOTS-c treatment (3x/week; 15mg/kg/day) would improve healthy lifespan (Fig. 3A). To assess healthspan, towards the end-of-life (>30 mo.), we performed a battery of physical tests to further probe the effect of MOTS-c on reversing age-dependent physical decline (Fig. 3A). LLII MOTS-c improved (i) grip strength (Fig. 4A), (ii) gait, assessed by stride length (Fig. 4B), and (iii) physical performance, assessed by a 60-second walking test (running was not possible at this age) (Fig. 4C). In humans, reduced stride length and walking capacity are strongly linked to mortality and morbidity (*36*). Together, these data indicate that LLII MOTS-c treatment improves physical capacity in old mice.

**Fig. 4.**
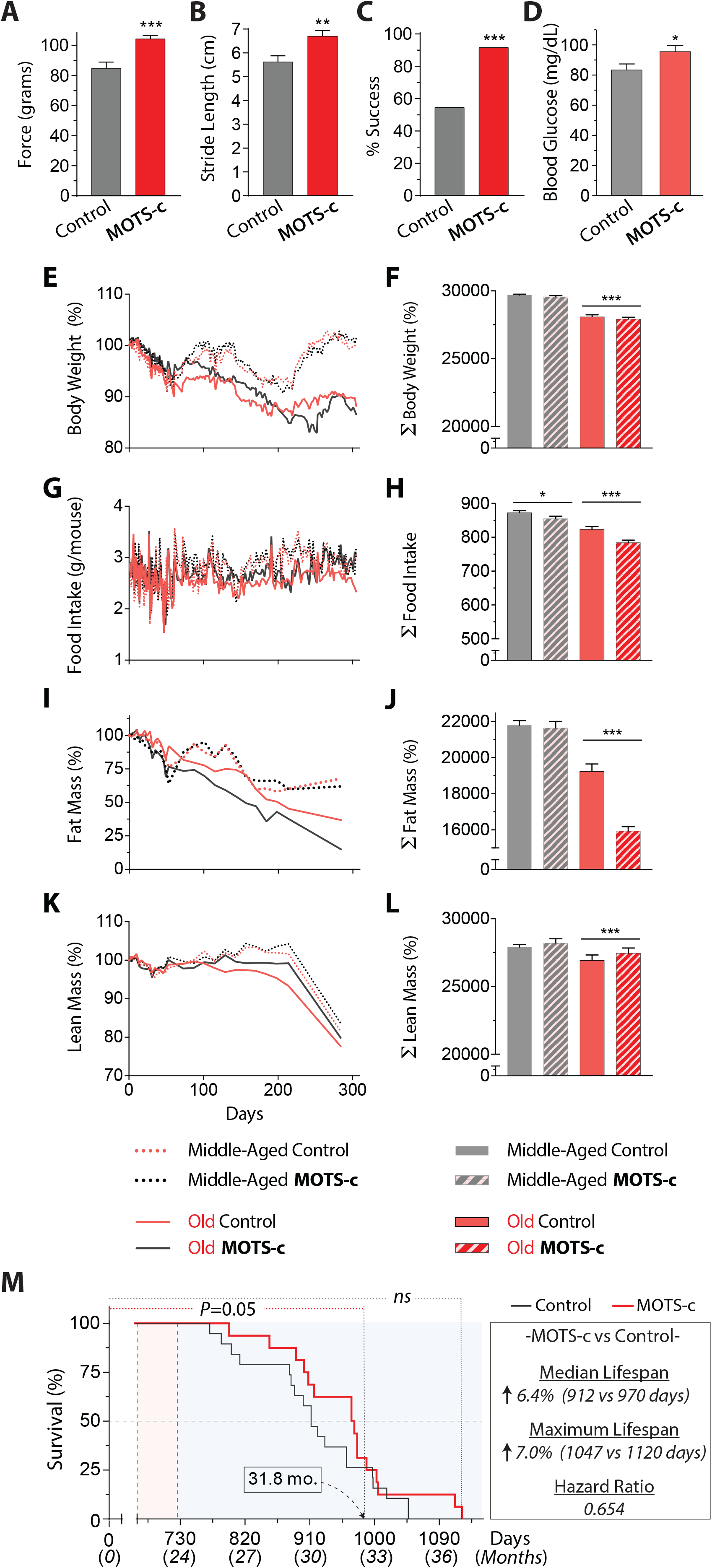
MOTS-c regulates aging metabolism and healthspan. Life-long measurements on male C57BL6/N mice treated intermittently (3x/week) with MOTS-c (15 mg/kg/day) starting at middle and old age (13.5 and 23.5 mo.) as described in Fig. 2a (n=16-19). (**A**) grip strength test (n=11), (**B**) gait analysis (stride length) (n=5), (**C**) 60-second walking test (n=11-12), and (**D**) blood glucose levels (n=11). (**E**, **F**) Body weight (**E**) as a function of time and (**F**) the total sum (∑); (**G**, **H**) Food intake (**G**) as a function of time and (**H**) the total sum (∑); (**I**, **J)** Percent fat mass (**I**) as a function of time and (**J**) the total sum (∑); (**K**, **L**) Percent lean mass (**K**) as a function of time and (**L**) the total sum (∑). (**M**) Lifespan curve; *P*=0.05 until 31.8 months of age. Overall curve trended towards increased median and maximum lifespan (*P*=0.23). Data expressed as mean +/− SEM. Log-rank (Mantel-Cox) test was used for (**M**). Otherwise, all statistics were performed using the Student’s *t*-test. \**P*<0.05, ** *P*<0.01, *** *P*<0.001.

Independent lines of research have shown that MOTS-c is a mitochondrial-encoded metabolic regulator at the cellular and organismal level (*13, 15, 24, 37–41*). We posited that LLII MOTS-c treatment would cause metabolic reprogramming in old mice. Consistent with our previous report (*13*), non-fasting blood glucose was better maintained in LLII MOTS-c-treated old mice (30 mo.; Fig. 4D). Over course of their life, LLII MOTS-c-treated mice showed comparable body weight to their untreated counterparts (Fig. 4E, F) However, total food intake was significantly reduced (Fig. 4G, H, fig. S6), whereas total activity was significantly higher (fig. S10). Body composition analysis using a time-domain NMR analyzer revealed significant reduction of fat mass (Fig. 4I, J) and a modest increase in lean mass (Fig. 4K, L). The RER, measured using metabolic cages, at 30 mo. revealed increased fat utilization, consistent with that obtained at ~23.5 mo. (Fig. 3F), but with a circadian shift (fig. S11); this is also consistent with reduced total fat mass (Fig. 4I, J, fig. S5B, E) and increased lipid utilization (*13, 41*). Ultimately, LLII MOTS-c treatment showed a trend towards increased median (6.4%) and maximum (7.0%) lifespan and reduced hazard ratio (0.654); *P*=0.05 until 31.8 months (Fig. 4M). Larger cohorts will be needed to confirm the broader significance of MOTS-c treatment on overall longevity. These data suggest that LLII MOTS-c treatment improves overall physical capacity in old mice and may compress morbidity and increase healthspan.

Skeletal muscle must adapt to various exercise-induced challenges (*42*), including nutrient (*e.g.* metabolic supply-demand imbalance) (*43*), oxidative (*44, 45*), and heat stress (*42, 46*), which share mitochondria as a common denominator. Because MOTS-c enhanced cellular resistance against metabolic/oxidative stress (*15*), we tested if MOTS-c treatment improved skeletal muscle adaptation to metabolic stress using C2C12 mouse myoblast cells. Using crystal violet staining to determine cellular viability, we found that MOTS-c (10μM) treatment significantly protected C2C12 cells (~2-fold) from 48 hours of metabolic stress [glucose restriction (GR; 0.5 g/L) and serum deprivation (SD; 1% FBS)] (Fig. 5A). Next, we tested the replicative capacity of C2C12 cells following prolonged metabolic stress as a functional marker of protection. C2C12 cells were metabolically stressed (GR/SD) for one week with daily MOTS-c (10μM) treatment, then replenished with complete medium for 2 days and stained with crystal violet. MOTS-c-treated C2C12 cells showed significantly enhanced proliferative capacity within 2 days (~6-fold) (Fig. 5B). Because MOTS-c promotes fat utilization, which may underly its effect on “metabolic flexibility” (Figs. 3F, 4I, J, fig. S5) (*13, 41, 47*), we tested if MOTS-c-treated C2C12 cells could survive on lipids without glucose (0 g/L). As expected, most control cells died without glucose even with lipid supplementation, whereas MOTS-c treatment provided significant protection (~2-fold) (Fig. 5C). Real-time metabolic flux analysis revealed that MOTS-c treatment significantly increased lipid utilization capacity (Fig. 5D) and lipid-dependent glycolysis (fig. S12) in C2C12 cells.

**Fig. 5.**
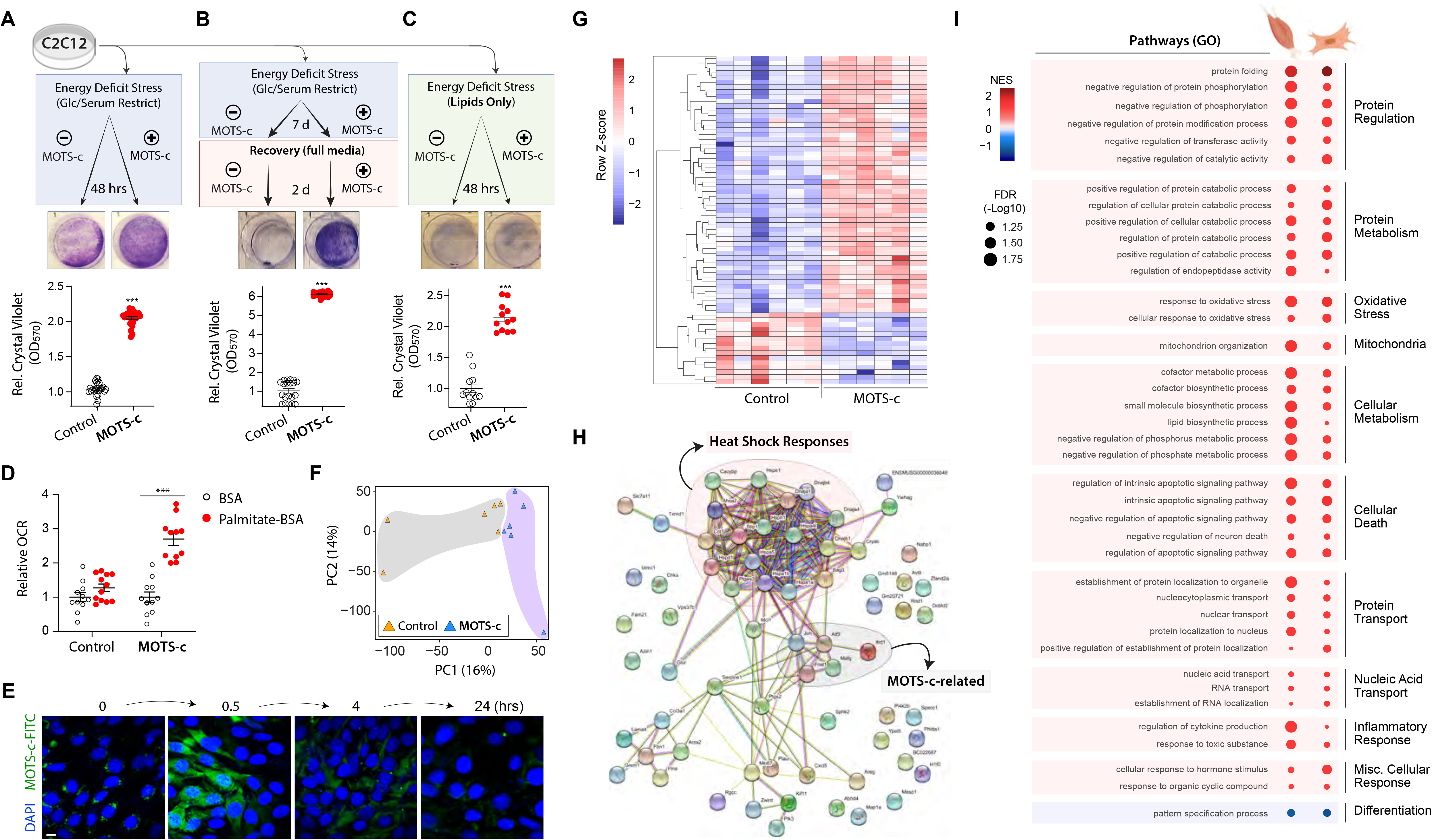
MOTS-c regulates myoblast gene expression and enhances adaptation to metabolic stress. (**A-C**) Survival of MOTS-c-treated (10μM; equal-volume vehicle control) C2C12 myoblasts assessed by crystal violet staining following (**A**) 48 hours of glucose restriction (GR; 0.5 g/L) and serum deprivation (SD; 1% FBS) with MOTS-c treated only once initially (n=12), (**B**) 7 days of GR/SD with daily MOTS-c treatment, followed by a 2-day recovery in full media with MOTS-c (n=10), and (**C**) 48 hours of complete GR (0 g/L) with chemically-defined lipid supplementation and daily MOTS-c treatment (n=6). (**D)** Real-time oxygen consumption rate (OCR) in response to fatty acid (palmitate-BSA) in C2C12 myoblasts treated with MOTS-c (10μM) for 48 hours (n=11-12). (**E**) Time-dependent subcellular localization pattern of exogenously treated MOTS-c-FITC (10μM) in C2C12 myoblasts. Scale bar: 10μm. (**F-I**) RNA-seq was performed on C2C12 myoblasts following 48 hours of GR/SD with/without a unique initial MOTS-c (10μM) treatment (n=6). (**F**) Principle Component Analysis (PCA) and (**G**) heatmap of significantly differentially regulated genes by MOTS-c at false discovery rate (FDR) < 5% by DESeq2 analysis. (**H**) Protein-protein interaction network analysis based on genes that were significantly differentially regulated by MOTS-c (FDR < 5%) using the STRING (Search Tool for the Retrieval of Interacting Genes/Proteins) database version 11.0 (*61*). (**I**) Balloon plots of common biological processes derived from RNA-seq data between MOTS-c-treated (i) skeletal muscle from old mice (see Fig. 2) and (ii) C2C12 myoblasts, based on gene set enrichment analysis (GSEA) using gene ontology biological process (GO_BP) (select gene sets; FDR < 15%). Data expressed as mean +/− SEM. Student’s *t*-test. \**P*<0.05, ** *P*<0.01, *** *P*<0.001

We previously reported that endogenous MOTS-c translocates to the nucleus to directly regulate adaptive nuclear gene expression in response to cellular stress (*15*). Using fluorescently labeled MOTS-c peptide (MOTS-c-FITC), we confirmed that exogenously treated MOTS-c also dynamically translocated to the nucleus in a time-dependent manner (Fig. 4E) (*15*), indicating a direct nuclear role. We performed RNA-seq on C2C12 cells treated with MOTS-c or vehicle control (10μM) under GR/SD for 48 hours and found (i) clustering by treatment type using a principal component analysis (PCA) (Fig. 5F) and (ii) 69 genes that were differentially regulated at FDR < 5% (Fig. 5G). Further, using the STRING (Search Tool for the Retrieval of Interacting Genes/Proteins) database to assess putative changes in protein-protein interaction networks based on our RNA-seq results, we found that a cluster related to heat-shock responses were prominently regulated by MOTS-c in C2C12 cells under GR/SD; we also identified previously reported MOTS-c targets, including Atf3, Jun, Fosl1, and Mafg (*15*) (Fig. 5H). Consistently, select GO_BP analysis revealed protein regulation as a major target process of MOTS-c in myoblasts (select terms in fig. S13; full results in table S3). To identify common pathways in *in vitro* and *in vivo* models, we overlaid RNA-seq data from MOTS-c-treated (i) old mouse skeletal muscle and (ii) metabolically stressed (GR/SD) C2C12. Select GO_BP analysis revealed several commonly targeted processes, including protein regulation, cellular metabolism, oxidative stress response, and nuclear transport (select terms in Fig. 5I; full results in table S4). Together, these data suggest that MOTS-c improves metabolic homeostasis/flexibility and protein homeostasis in skeletal muscle under exercise-induced stress conditions.

Our study shows that exercise induces mtDNA-encoded MOTS-c expression. MOTS-c treatment significantly (i) improved physical performance in young, middle-aged, and old mice, (ii) regulated skeletal muscle metabolism and gene expression, and (iii) enhanced adaptation to metabolic stress in C2C12 cells. Thus, it is plausible that the physiological role of exercise-induced MOTS-c is to promote adaptive responses to exercise-related stress conditions (*e.g.* metabolic imbalance and heat shock) in the skeletal muscle and maintain cellular homeostasis.

Mitochondria are strongly implicated in aging at multiple levels (*1, 2, 6–8, 48*). Here, we present evidence that the mitochondrial genome encodes for instructions to maintain physical capacity (*i.e.* performance and metabolism) during aging and thereby increase healthspan. MOTS-c treatment initiated in late-life, proximal to the age at which the lifespan curve rapidly descends for C57BL/6N mice, significantly delayed the onset of age-related physical disabilities, suggesting “compression of morbidity” in later life (*49*). Interestingly, an exceptionally long-lived Japanese population harbors a mitochondrial DNA (mtDNA) SNP (m.1382A>C) that yields a functional variant of MOTS-c (*50, 51*).

Our study shows that exogenously treated MOTS-c enters the nucleus and regulates nuclear gene expression, including those involved in heat shock response and metabolism. Thus, age-related gene networks are comprised of integrated factors encoded by both genomes, which entails a bi-genomic basis for the evolution of aging. Although the detailed molecular mechanism(s) underlying the functions of MOTS-c is an active field of research, we provide a “proof-of-principle” study that realizes the mitochondrial genome as a source for instructions that can regulate physical capacity and healthy aging.

## Materials and Methods

### Mouse Care

All animal work was performed in accordance with the University of Southern California (USC) Institutional Animal Care and Use Committee. MOTS-c (New England Peptide, USA) was administered daily at 5 or 15 mg/kg via intraperitoneal injections. 12-week old male CD-1 (outbred) mice (Charles River, USA), 12-week old male C57BL/6J mice (Jackson Laboratory) and 8- and 18-month old male C57BL/6N mice (National Institute on Aging; NIA) were obtained. All mice were fed either a HFD (60% calories from fat) or matching control diet (Research Diets, USA, #D12492 and D12450J, respectively). NIA mice were sufficiently acclimated for 4 months in our vivarium until they were considered middle-aged (12 mo.) and old (22 mo.) at the start of MOTS-c injections. Body weight and food consumption were recorded daily, while body composition was analyzed twice weekly using an LF90II time-domain NMR minispec (Bruker, USA). After eight weeks of injections (23.5 months of age), mice were transitioned to receive MOTS-c injections three times weekly. No live mouse was censored.

### Physical Tests in Mice

#### Running Test

Prior to running training/testing, mice were acclimated to the stationary treadmill apparatus (TSE-Systems, USA) for ten minutes on two consecutive days (Days 1 and 2). Both the high intensity test and training protocols were adapted from previously published protocols (*52*). Running training was given twice on non-consecutive days and consisted of a fixed speed run of 10 m/min for 20 minutes (Days 4 and 6) on a level treadmill. The treadmill test on Day 10 consisted of three stages. Stage one was a five-minute run at 13 m/min. For the next five minutes, the speed was increased by 1 m/min. The mice run at a fixed speed of 18 m/min for the next 30 minutes. Finally, after 40 minutes of total run time, the running speed is increased to 23 m/min until exhaustion is reached. All training and testing were done on a level treadmill. Mice resting on the platform were gently prodded to encourage re-engagement. Any mouse that resisted prodding and remained on the platform for 30 seconds was considered to be exhausted, and time was recorded.

#### Walking Test

When the mice reached 30 months of age, they were no longer capable of performing the same treadmill routine. We developed a measure of mobility in the aged mice consisting of a 60 second walking test. The treadmill was set at 13 m/min for 60 seconds. We recorded whether the mouse was able to walk, or not, on the treadmill for 60 seconds, with gentle prodding as needed. Mice remaining on the stationary platform, refusing to engage in the treadmill walking, for more than five seconds were considered to have failed the test.

#### Rotarod

The Rotarod test was performed by placing the mice on the apparatus (TSE-Systems), all facing the opposite direction of rotation. The initial speed of rotation was 24 rpm and accelerated at 1 rpm every 10 seconds. Time to fall was recorded for each mouse, and three trials per mouse was run. Mice received no less than five minutes of recovery time between trials.

#### Grip Strength

We measured grip strength using a horizontal bar connected to the grip strength meter (TSE-Systems) as a high precision force sensor for the forelimbs. After allowing the mouse to properly grip the bar they were firmly and quickly pulled in the opposite direction. Only trials where the mouse released its paws from the bar simultaneously were counted as successful. Mice underwent three trials, with at least 30 seconds recovery time between trials.

#### Gait Analysis

To perform gait analysis, we applied a different color of non-toxic ink (BLICK^®^, USA) to the front and hind paw of the mice to record footprints. Barriers were constructed to guide the mice to walk straight on the recording paper. The home cage was kept at the end of the recording paper to encourage completion of the test. Only trials in which the mouse made a continuous, direct path to its home cage were counted. Stride length was measured as the average forward movement of three full strides as previously described(*53*).

### Cognitive tests

*Y-maze tests* were performed as previously described (*54*). Briefly, mice were placed in a maze consisting of three arms equally spaced 120° apart. Mice were placed in one arm of the maze and allowed to freely explore the maze for five minutes. Total arm entries and arm choices were recorded for each mouse. An arm entry was defined as a mouse having both front and hind paws entering the arm fully. Percent alternations was defined as an arm choice differing from the previous two compared to the total number of alternation opportunities.

*Barnes maze* tests were performed as previously described (*54*). 12-week old male CD-1 mice were tested twice daily for 7 days. Mice were placed in a start chamber in the middle of the maze and allowed to habituate (30 seconds), then the mouse was released to explore the maze and find the escape box (EB). Latency (time to enter the EB) and number of errors (nose pokes and head deflections over false holes) were recorded. A maximum of 2 minutes was allowed for each trial.

### *In Vivo* Metabolism Assessment

#### Metabolic Cages

Metabolic activity in mice was measured using the PhenoMaster system (TSE-Systems) equipped to detect indirect calorimetry, measure food and water intake, and monitor activity. Prior to metabolic analysis, mice were housed 3-4 per cage in a facility with a 12:12 hour light-dark cycle (light period 0600-1800) at 24°C. Food and water were available *ad libitum*. For metabolic assessment, nice were moved into individual PhenoMaster cages in an isolated room under the same environmental conditions. Mice were automatedly monitored for 36 hours to record physiological parameters. To measure O_2_ intake and CO_2_ production, gas sensors were calibrated prior to the study using primary gas standards of known concentrations of O_2_, CO_2_, and N_2_. Room air was passed through the animal chambers at a rate of 0.5 L/min. Exhaust air from individual cages were sampled at 30-minute intervals for 3 minutes. Sample air was passed through sensors to determine oxygen consumption (VO_2_) and carbon dioxide production (VCO_2_). The respiratory exchange ratio (RER) was calculated as the ratio of carbon dioxide produced to oxygen consumption. The PhenoMaster system allows for activity monitoring using a triple beam IR technology system. Breaking the IR beams through movement was considered a “count”. The three-beam system allows XYZ monitoring that considers both ambulatory activity around the cage as well as rearing activity. All data are expressed as the mean of three 24-hour acquisition cycles.

#### Blood glucose

Blood was collected via a single tail-nick and immediately analyzed using a glucometer (Freestyle, Abbott). Blood collection was performed by trained professionals and in accordance with the University of Southern California Institutional Animal Care and Use Committee.

### Western Blots

Protein samples were lysed in 1% Triton X-100 (Thermo Fisher Scientific, USA, #21568-2500) with 1 mM EDTA (Promega Life Sciences, USA, #V4231) and 100 mM Tris-HCl pH 7.5 (Quality Biological, USA, #351-006-101) and protease inhibitors (Roche, Germany, #118636170001) and sonicated using a Sonic Dismembrator (Fisher Scientific, USA). Samples were heated at 95°C for five minutes. Samples were ran on 4-20% gradient tris-glycine gels (TGX; Bio-Rad, USA, #456-1104) and transferred onto 0.2 μM PVDF membranes (Bio-Rad #162-0184) using a Transblot Turbo semi-dry transfer system (Bio-Rad) at 9 volts for 15 minutes. Membranes were blocked for 1 hour using 5% BSA (Akron Biotech, USA, #AK8905-0100) in tris-buffered saline containing 0.05% Tween-20 (Bio-Rad #161-0781) and incubated in primary antibodies against MOTS-c (rabbit polyclonal; YenZym, USA) and GAPDH (cat# 5174; Cell Signaling, USA) overnight at 4°C. Secondary HRP-conjugated antibodies (#7074; Cell Signaling, USA) were then added (1:30,000) for one hour at room temperature. Chemiluminescence was detected and imaged using Clarity western ECL substrate (Bio-Rad #1705060) and Chemidoc XRS system (Bio-Rad). Western blots were quantified using ImageJ version 1.52k.

### Cell Studies

#### Cell culture

C2C12 cells were cultured in DMEM with 4.5 g/L glucose (Corning, USA #10-017-CV) and 10% FBS (Millipore-Sigma, USA, #F0926-500). All cells were stored at 37°C and 5% CO_2_. Cells were passaged when they reached 75-80% confluence using TrypLE (Thermo Fisher Scientific #12605-010).

#### Cell survival assays

Protection against glucose restriction (GR) and serum deprivation (SD) was tested by culturing cells in DMEM (Thermo Fisher Scientific #11966-025) with 0.5 g/L glucose (Millipore-Sigma #G8769) and 1% FBS. MOTS-c (10μM) or vehicle (PBS). MOTS-c (10μM) was added to the media every 24 hours. After 48 hours of GR/SD, we performed crystal violet (Thermo Fisher Scientific #C581-25) staining as before (*15*) to determine cell survival. We also tested cellular proliferation, following prolonged (7-day) GR/SD (DMEM with 1% FBS and 0.5 g/L glucose), as a measure of cellular fitness. In this case, MOTS-c-containing (10μM) media was changed once every two days; no additional MOTS-c supplementation was given between media changes. After 7 days of GR/SD, we returned the cells to full growth media (10% FBS and 4.5 g/L glucose) for 48 hours with MOTS-c (10μM), then stained them with crystal violet. To determine the metabolic flexibility to utilize fatty acids, we cultured cells in DMEM with 1% FBS, 0.5 g/L glucose, and 1% chemically defined lipid mixture (Millipore-Sigma #L0288) for 48 hours, then stained them with crystal violet.

#### Metabolic flux

Real-time oxygen consumption and extracellular acidification rates in C2C12 myoblasts treated with 16% palmitate-BSA (1mM palmitate conjugated to 0.17mM BSA) or 16% BSA (0.17 mM; Seahorse Bioscience #102720-100) were obtained using the XF96 Bioanalyzer (Seahorse Bioscience) at the USC Leonard Davis School of Gerontology Seahorse Core. All values were normalized to relative protein concentration using a BCA protein assay kit (Thermo Fisher Scientific #23227).

#### Confocal microscopy

Confocal images were obtained using a Zeiss Confocal Laser Scanning Microscope 700 (Zeiss, Germany). C2C12 myoblasts were cultured on glass coverslips (Chemglass, USA, #CLS-1760-015). Cells were treated with FITC-MOTS-c (New England Peptide) for either 0 hours (immediate), 30 minutes, 4 hours, or 24 hours. Some cells were left as untreated controls (data not shown). All cells were treated with Hoeschst (Biotium, USA, #40045) for 15 minutes and then washed three times with PBS. Cells were fixed in 10% formalin (Millipore-Sigma #EM-R04586-82) and washed an additional three times in PBS. Coverslips were affixed to glass slides (VWR, USA, #48300-025) using ProLong Gold antifade reagent (Life Technologies Corporation, USA, #P36934).

### Human Studies

#### Study outline

Participants gave written consent before the commencement of the study, which was approved by the Northern Health and Disability Ethics Committee (New Zealand) (16/STH/116/AM01). 10 sedentary (<4h aerobic exercise/week) healthy young males (24.5 ± 3.7 years old and BMI 24.1 ± 2.1) were recruited to take part in a two-visit exercise trial. Recruited participants were free of cardiovascular, metabolic and blood diseases and were not taking any medication or supplements. The trial was separated into two visits, each involving exercise bouts that were carried out on an electromagnetically braked cycle ergometer (Velotron, RacerMate, USA).

##### Determination of peak oxygen uptake (VO_2_peak) and maximal power output (visit 1)

Peak oxygen uptake was determined using a ramped cycling exercise protocol. Prior to testing, participants warmed up for five minutes at a self-selected workload between 60 and 80W. The ramp protocol began at 60W, with the cycling power output set to increase by 1W every 4 seconds (15W/min) continuously until the participant was unable to maintain cycling workload (cycling cadence<minimum 60 revolutions per minute) or maximal volitional fatigue was reached. Mean Peak oxygen uptake of participants was reported as 38.4 ± 7.3 ml.kg.min.

##### Acute high intensity cycling exercise session (visit 2)

Prior to visit 2, participants were asked to fast overnight (from 10PM) and were instructed to abstain from physical activity for at least 48h prior. Upon arrival to the laboratory, participants lay supine for 15-minutes and then had an intravenous cannula inserted into a forearm vein. A resting plasma sample was collected followed by a pre-exercise muscle biopsy taken from the vastus lateralis muscle (quadriceps muscle). Approximately 10-minutes later, participants completed ten, 60-second cycling intervals at individually-specified peak power workloads (determined from peak oxygen uptake test) followed by 75-seconds of rest/low intensity cycling (<30W) per interval as previously described (*55*). A mid-exercise blood sample was taken following the completion of the 5^th^ exercise interval as well as immediately following the completion of the exercise bout. In addition, an immediately-post exercise muscle biopsy was taken (within ~5-minutes of completion of the exercise bout). Participants remained supine and resting in the procedure bed for a 4-hour recovery period. Following four-hours of recovery, a final blood and muscle biopsy sample was collected.

##### Muscle biopsy and blood sampling

Muscle biopsies were extracted under local anesthesia (1% xylocaine) using the Bergstrom needle with manual suction technique(*56*). Biopsies were snap-frozen in liquid nitrogen and stored at −80°C until analyzed. Blood was drawn through a 20-gauge cannula, collected in 10-mL EDTA vacutainers and then centrifuged immediately upon collection at 4°C at 2,000 g for 10 minutes. Plasma was extracted and then stored at −80°C until further analysis using an in-house ELISA as described before (*13*). Human skeletal muscle was processed for western blotting by soaking the samples in lysis buffer (above) and minced using a razor blade. Once the sample was evenly minced, we proceeded with the Sonic Dismembrator step as described above.

### Liquid Chromatography-Mass Spectrometry Metabolomics

Metabolites were extracted from randomly selected tissue samples by adding 1 mL of 80:20 methanol:water solution on dry ice. Samples were incubated at −80C for 4 hours and centrifuged at 4C for 5 minutes at 15k rpm. Supernatants were transferred into LoBind Eppendorf microcentrifuge tubes and the cell pellets were re-extracted with 200 μL ice-cold 80% MeOH, spun down and the supernatants were combined. Metabolites were dried at room temperature under vacuum and re-suspended in water for injection.

Samples were randomized and analyzed on a Q-Exactive Plus hybrid quadrupole-Orbitrap mass spectrometer coupled to an UltiMate 3000 UHPLC system (Thermo Scientific). The mass spectrometer was run in polarity switching mode (+3.00 kV/-2.25 kV) with an m/z window ranging from 65 to 975. Mobile phase A was 5 mM NH4AcO, pH 9.9, and mobile phase B was acetonitrile. Metabolites were separated on a Luna 3 μm NH2 100 Å (150 × 2.0 mm) column (Phenomenex). The flowrate was 300 μl/min, and the gradient was from 15% A to 95% A in 18 min, followed by an isocratic step for 9 min and re-equilibration for 7 min. All samples were injected twice for technical duplicates. Metabolites were detected and quantified as area under the curve based on retention time and accurate mass (≤ 5 ppm) using the TraceFinder 3.3 (Thermo Scientific) software.

### RNA-seq

#### RNA purification from tissue and cells

Total RNA extraction from skelatal muscle tissue or C2C12 mouse myoblasts was done using TRI Reagent (Millipore-Sigma #T9424). Muscle tissue samples were flash-frozen in liquid nitrogen until further processing. Tissues were resuspended in 600μL of TRI Reagent, then homogenized on Lysing Matrix D 2mL tubes (MP Biomedicals) on a BeadBug homogenizer (Benchmark Scientific). For both skeletal muscle and C2C12 cells, total RNA was purified using the Direct-zol RNA MiniPrep (Zymo Research #R2052).

#### RNA-seq library preparation

Total RNA was subjected to rRNA depletion using the NEBNext rRNA Depletion Kit (New England Biolabs), according to the manufacturer’s protocol. Strand specific RNA-seq libraries were then constructed using the SMARTer Stranded RNA-Seq Kit (Clontech # 634839), according to the manufacturer’s protocol. Based on rRNA-depleted input amount, 13-15 cycles of amplification were performed to generate RNA-seq libraries. Paired-end 150bp reads were sent for sequencing on the Illumina HiSeq-Xten platform at the Novogene Corporation (USA). The raw sequencing data was deposited to the NCBI Sequence Read Archive (accession: PRJNA556045). The resulting data was then analyzed with a standardized RNA-seq data analysis pipeline (described below).

#### RNA-seq analysis pipeline

To avoid the mapping issues due to overlapping sequence segments in paired end reads, reads were hard trimmed to 75bp using the Fastx toolkit v0.0.13. Reads were then further quality-trimmed using Trimgalore 0.4.4 (github.com/FelixKrueger/TrimGalore) to retain high-quality bases with Phred score > 20. All reads were also trimmed by 6 bp from their 5’ end to avoid poor qualities or biases. cDNA sequences of protein coding and lincRNA genes were obtained through ENSEMBL Biomart for the GRCm38 build of the mouse genome (Ensemble release v94). Trimmed reads were mapped to this reference using kallisto 0.43.0-1 and the –fr-stranded option (*57*). All subsequent analyses were performed in the R statistical software (https://cran.r-project.org/).

Read counts were imported into R, and summarized at the gene level, to estimate differential gene expression as a function of age.

Because of high sample variability, we used surrogate variable analysis to remove experimental noise from the muscle RNA-seq dataset (*58*). R package ‘sva’ v3.24.4 (*59*) was used to estimate surrogate variable, and the effects of surrogate variables were regressed out using ‘limma’. Corrected read counts were then used for downstream analyses.

DEseq2 normalized fold-changes were then used to estimate differential gene expression between control and MOTS-c treated muscle or cell samples using the ‘DESeq2’ R package (DESeq2 1.16.1)(*60*). The heatmap of expression across samples for significant genes (Fig. 4G) was plotted using the R package ‘pheatmap’ 1.0.10 (Raivo Kolde, 2015-12-11; https://CRAN.R-project.org/package=pheatmap). Putative protein-protein interaction was derived using the STRING (Search Tool for the Retrieval of Interacting Genes/Proteins) database version 11.0 (*61*) (https://string-db.org/).

### Functional enrichment analysis

To perform functional enrichment analysis, we used the Gene Set Enrichment Analysis paradigm through its implementation in the R package ‘ClusterProfiler’ v3.10.1 (*62*), and Bioconductor annotation package ‘org.Mm.eg.db’ v3.7.0. Balloon plots representing the output were generated using R packages ‘ggplot2’ v3.1.0 and ‘scales’ 1.0.0.

### Principal Component Analysis

#### Metabolites

Principal component analysis (PCA) was performed using the mean-centered matrix of metabolite values per each mouse. Principal components that separated sample groups were identified with visual inspection. Loadings from principal components that stratify experimental samples versus controls were then queried against metabolic pathways using a Kolmogorov-Smirnov statistic against the expected distribution of metabolites. Metabolic pathway enrichment analysis (gene set enrichment analysis, GSEA) (*63*) were performed using 28 metabolic pathways defined by the Kyoto Encyclopedia of Genes and Genomes (KEGG) database using pathways with four or more measured metabolites.

#### RNA-seq

PCA was performed using the R base package function ‘prcomp’. The first 2 principal components were used.

### Quantification and Statistical Analysis

Unless otherwise noted, statistical significance was determined using the Student *t*-test. Statistical tests were performed using GraphPad Prism version 8.1.2. Results of *t*-tests are indicated in all figures as \**p*<0.05, ** *p*<0.01, \*\*\**p* <0.001 and *ns* for not significant (*p*>0.05).

The RNA-seq analytical code will be made available on the Benayoun lab github (https://github.com/BenayounLaboratory/MOTSc_Exercise).

## Acknowledgments

We thank the USC Leonard Davis School of Gerontology Mouse Phenotyping core and Seahorse Bioanalyzer core and the USC Genomics core for experimental assistance. Funding was provided by the American Federation for Aging Research (AFAR), the National Institute on Aging (T32 AG052374) and the USC Manning Endowed Fellowship to J.C.R., a Mork Graduate Fellowship from the USC Viterbi School of Engineering to J.H.J., the USC Viterbi School of Engineering to N.A.G., the NIA (P01AG034906) to P.C., the NIA (R00AG049934), an innovator grant from the Rose Hills foundation, a seed grant the NAVIGAGE foundation, and the Hanson-Thorell Family to B.A.B., Rutherford Discovery Fellowship and a Marsden Fund Fast Start Grant to T.L.M., and the NIA (R01AG052258), Ellison Medical Foundation (EMF), AFAR, and the Hanson-Thorell Family to C.L.

## Author contributions

J.C.R., T.L.M., B.A.B., P.C., N.A.G., D.C-S., and C.L. conceived the experiments. J.C.R., J.S.T.W., B.A.B., R.L., R.W.L., J.H.J., C.J.M., and D.C-S. performed experiments. J.C.R., T.L.M., J.S.T.W., B.A.B., R.L., R.W.L., P.C., N.A.G., J.H.J., and C.L. analyzed the data. J.C.R. and C.L. wrote the manuscript.

## Data and materials availability

All data are available in the main manuscript and Extended Data material. RNA-seq data have been uploaded to the NCBI SRA database (accession: PRJNA556045).

---

**Fig. S1.**
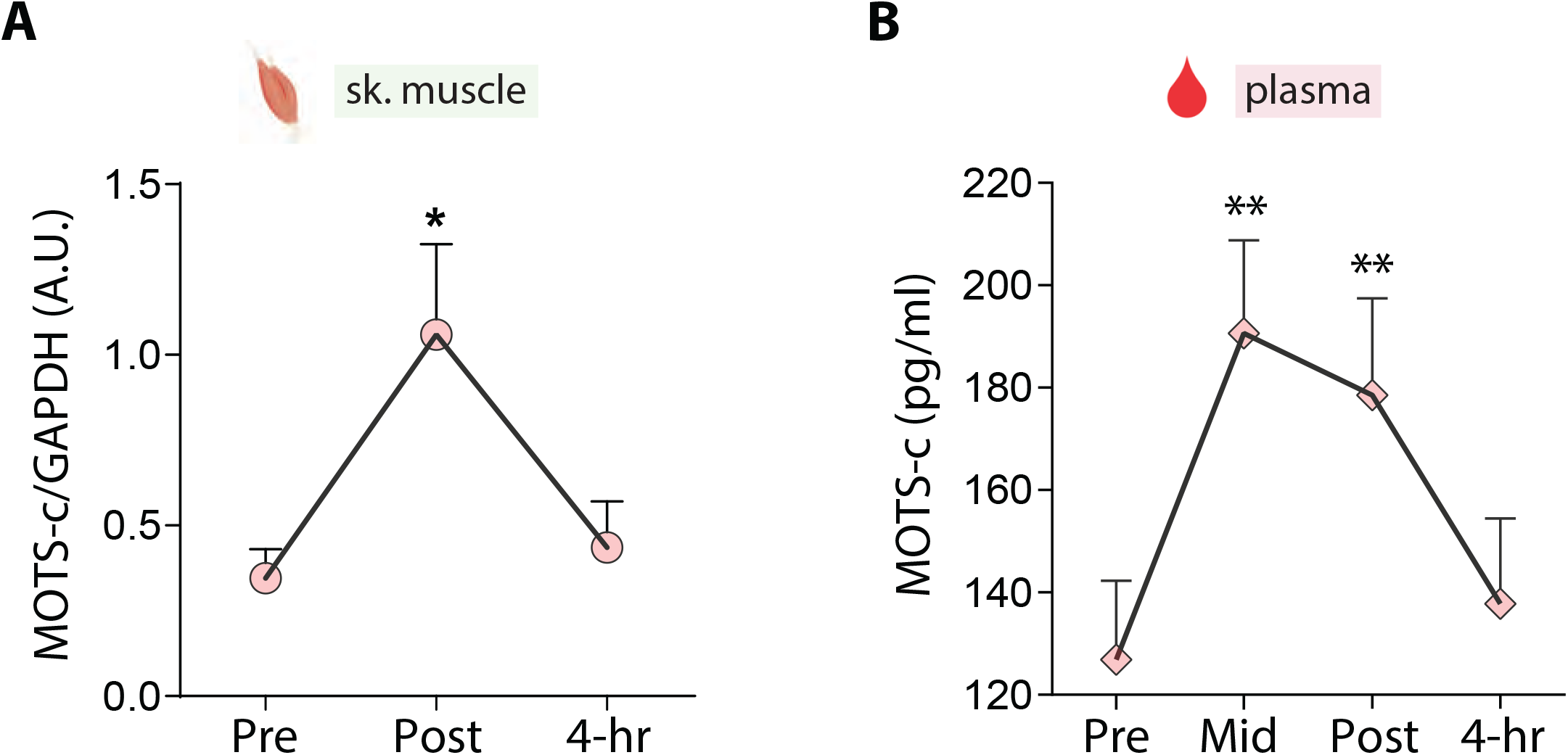
MOTS-c levels in human muscle and plasma. MOTS-c levels measured by (A) Western blotting on human skeletal muscle collected pre-, post-exercise and 4-hours of resting and (B) ELISA on plasma from same individuals collected pre-, mid-, post-exercise and 4-hours of resting (n=10). Statistics by Wilcox-on matched-pairs signed rank test. **P<0.01

**Fig. S2.**
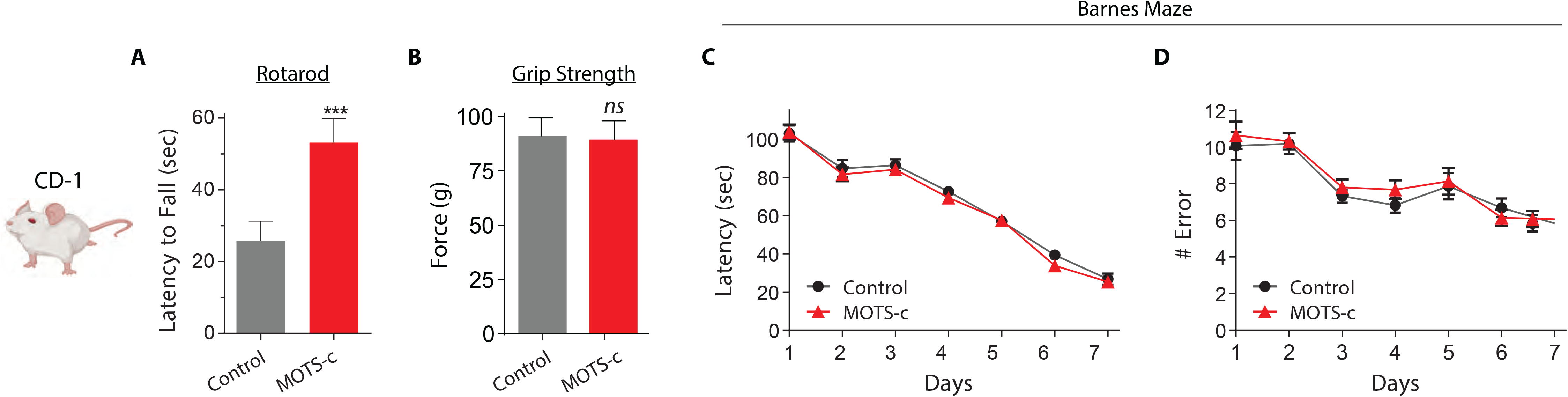
Rotarod, grip strength, and Barnes Maze tests in MOTS-c treated old mice. (**A**) Summary of latency time to fall on the Rotarod test (n=15). The speed of the rotations increased from a starting speed of 24 rpm by 1 rpm every 10 seconds. (**B**) grip strength test. (**C**,**D**) Barned Maze performance in control and MOTS-c treated 12-week old CD-1 mice (n=15). (**C**) There was no changes in average time to find the escape box (latency) between control and MOTS-c treated mice. (**D**) There was no change in the number of errors made prior to discovering the escape box between groups. Errors were defined as nose-pokes or head deflections over false holes. Data expressed as mean +/− SEM of three 24-hour acquisition cycles. Student’s t-test. *P<0.05, **P<0.01, ***P<0.001.

**Fig. S3.**
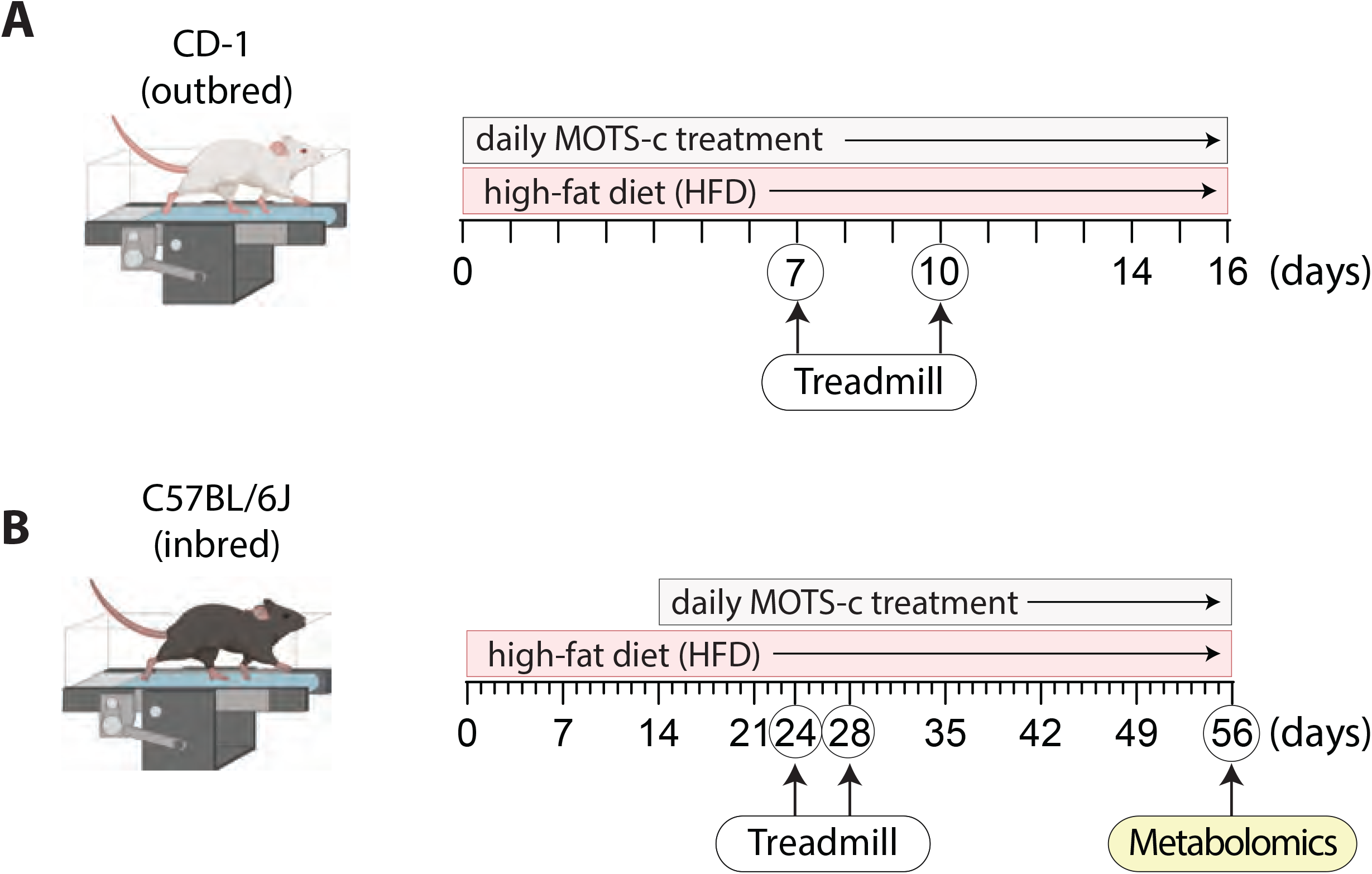
Outline of HFD mouse experiments. Timeline of experiment for 12-week old male CD-1 (outbred) and C57BL/6J (inbred) mice fed a HFD or defined control diet. (**A**) CD-1 mice were fed a HFD and given daily intraperitoneal injections (IP) of MOTS-c (0, 5, or 15 mg/kg/day) from Day 0. Treadmill running tests were performed on Day 7 (fig. S4a) and Day 10 (Fig. 1e-h). Daily MOTS-c injections ceased at Day 16. (**B**) C57BL/6J mice were started on either a HFD or a defined control diet on Day 0 and continued uninterruptedly throughout the experiment. Daily MOTS-c treatment (15 mg/kg; IP) started on Day 14. Treadmill running tests were performed on Day 24 and Day 28 (10 days and 14 days after the start of MOTS-c treatment) (Fig. 1i-l; fig. S4b). Mice were treated daily until Day 56, at which time metabolomics was performed (Fig. 1m).

**Fig. S4.**
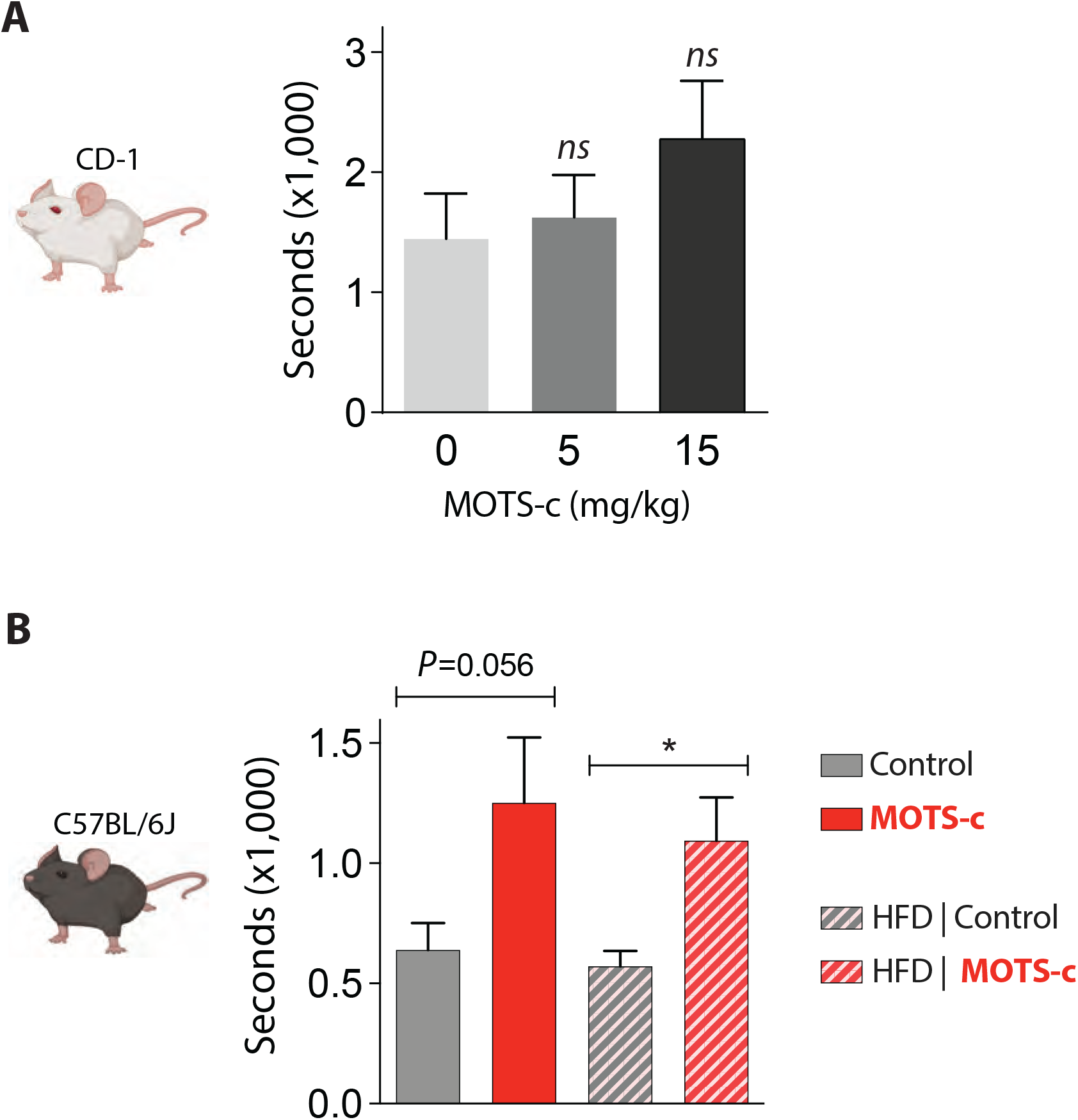
Initial running time of MOTS-c-treated young mice. (**A**) Running time of CD-1 mice following seven days of MOTS-c treatment (n=5-6). MOTS-c (15 mg/kg/day) treatment showed a trend towards enhanced running performance. (**B**) Running time of HFD-fed C57BL/6J mice following 10 days of MOTS-c treatment (n=8). Data expressed as mean +/− SEM. Student’s t-test. *P<0.05

**Fig. S5.**
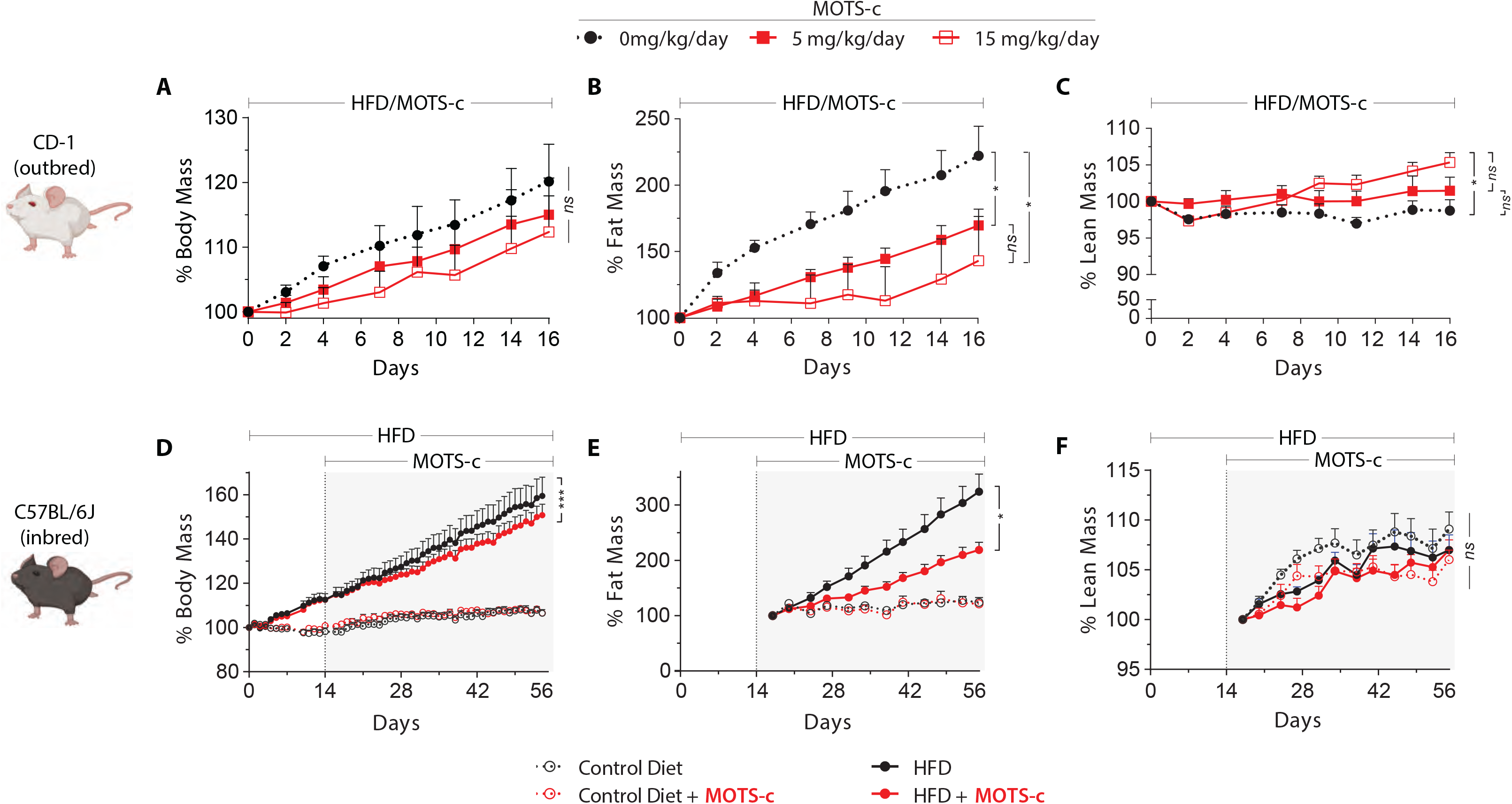
Body composition analysis on MOTS-c-treated young mice. Body composition was measured non-invasively using a time-domain NMR analyzer. (**A-C**) Young CD-1 mice were treated daily with MOTS-c (0, 5, or 15 mg/kg/day;IP) for 16 days (n=5-6) and percent (**A**) body weight, (**B**) fat mass, and (**C**) lean muscle mass were measured. (**D-F**) C57BL/6J mice either on a HFD or a defined control diet and treated daily with MOTS-c (15 mg/kg/day; IP) or saline control (n=8) and percent (**D**) body weight, (**E**) fat mass, and (**F**) lean muscle mass were measured. The dotted line at Day 14 represents the start of MOTS-c treatment. Data expressed as mean +/− SEM. Significance determined by using two-way ANOVA (repeated measures). *P<0.05, **P<0.01, ***P<0.001.

**Fig. S6.**
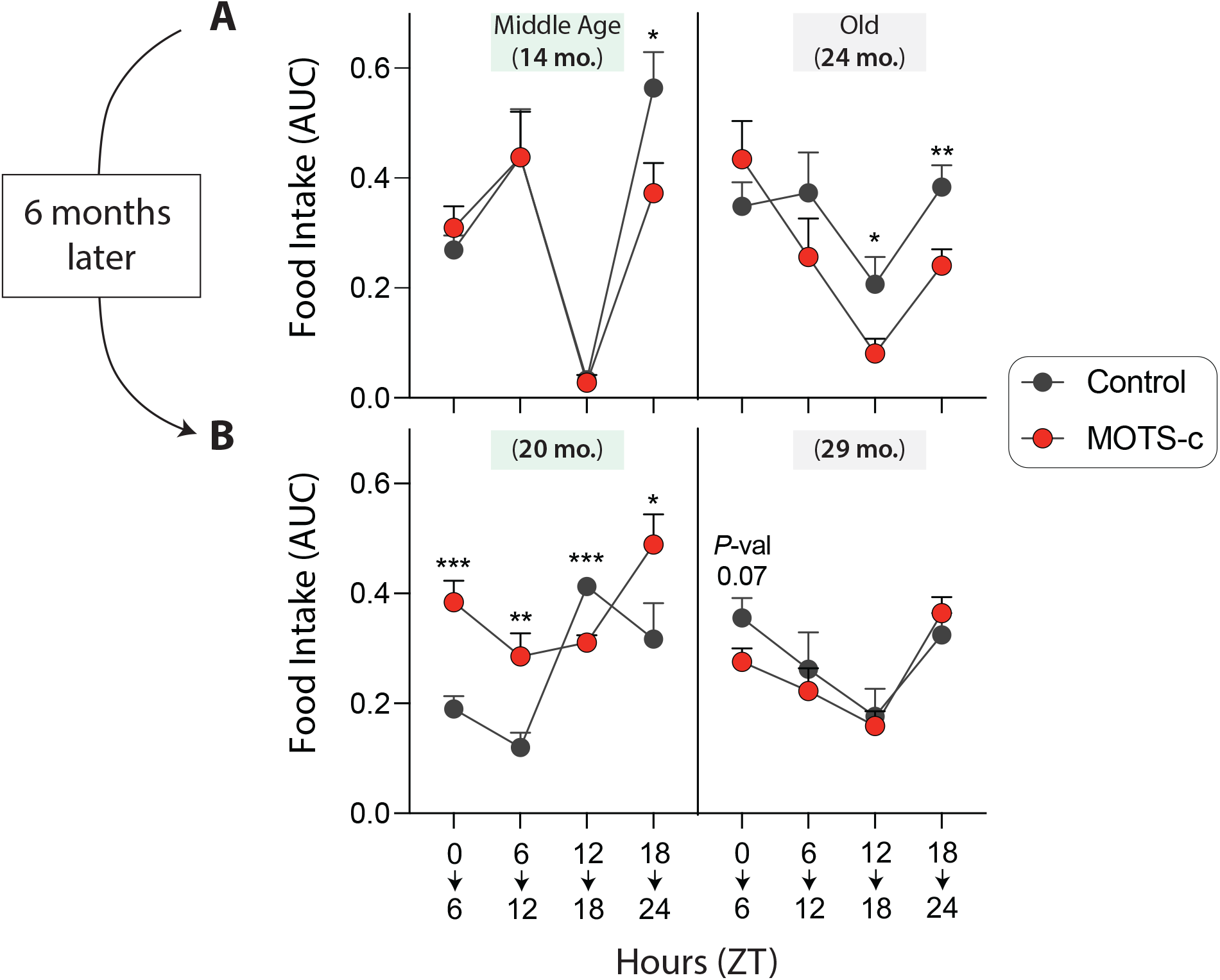
Circadian pattern of food intake in MOTS-c-treated old mice. (**A**) The sum of continuous food intake measurements using metabolic cages divided into daily quartiles in MOTS-c-treated middle-age (14 months) and old (24 months) mice (n=4). (**B**) Measurements were taken 6 month later in same mice. Data expressed as mean +/− SEM. Student’s t-test. *P<0.05, **P<0.01, ***P<0.001.

**Fig. S7.**
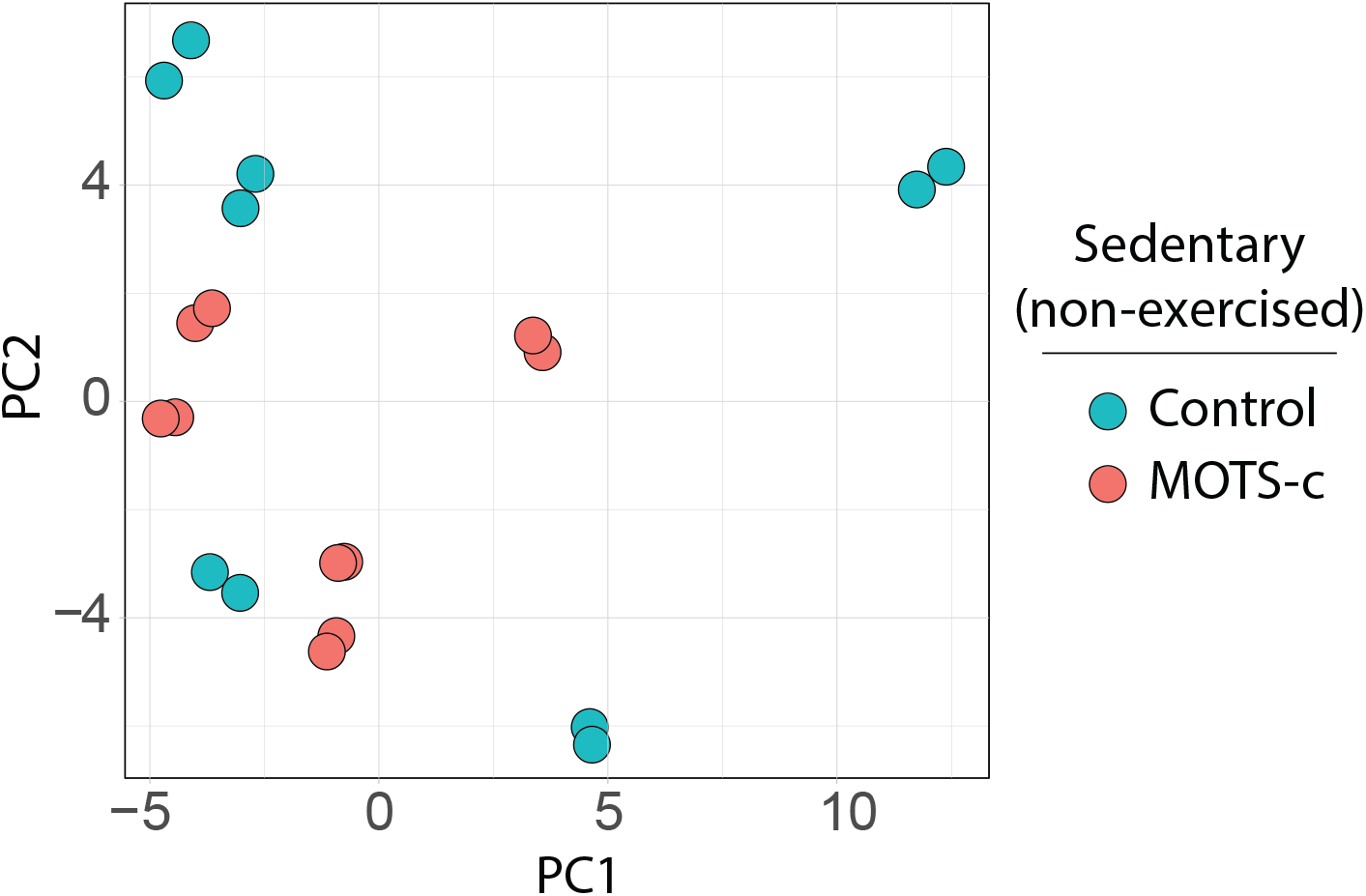
Metabolomic analysis on sedentary MOTS-c-treated old mice. Skeletal muscle from sedentary (not treadmill-exercised) old mice (22.5 months) treated daily with MOTS-c (15 mg/kg/day) for 2 weeks (n=10) were subject to metabolomics and analyzed using PCA.

**Fig. S8.**
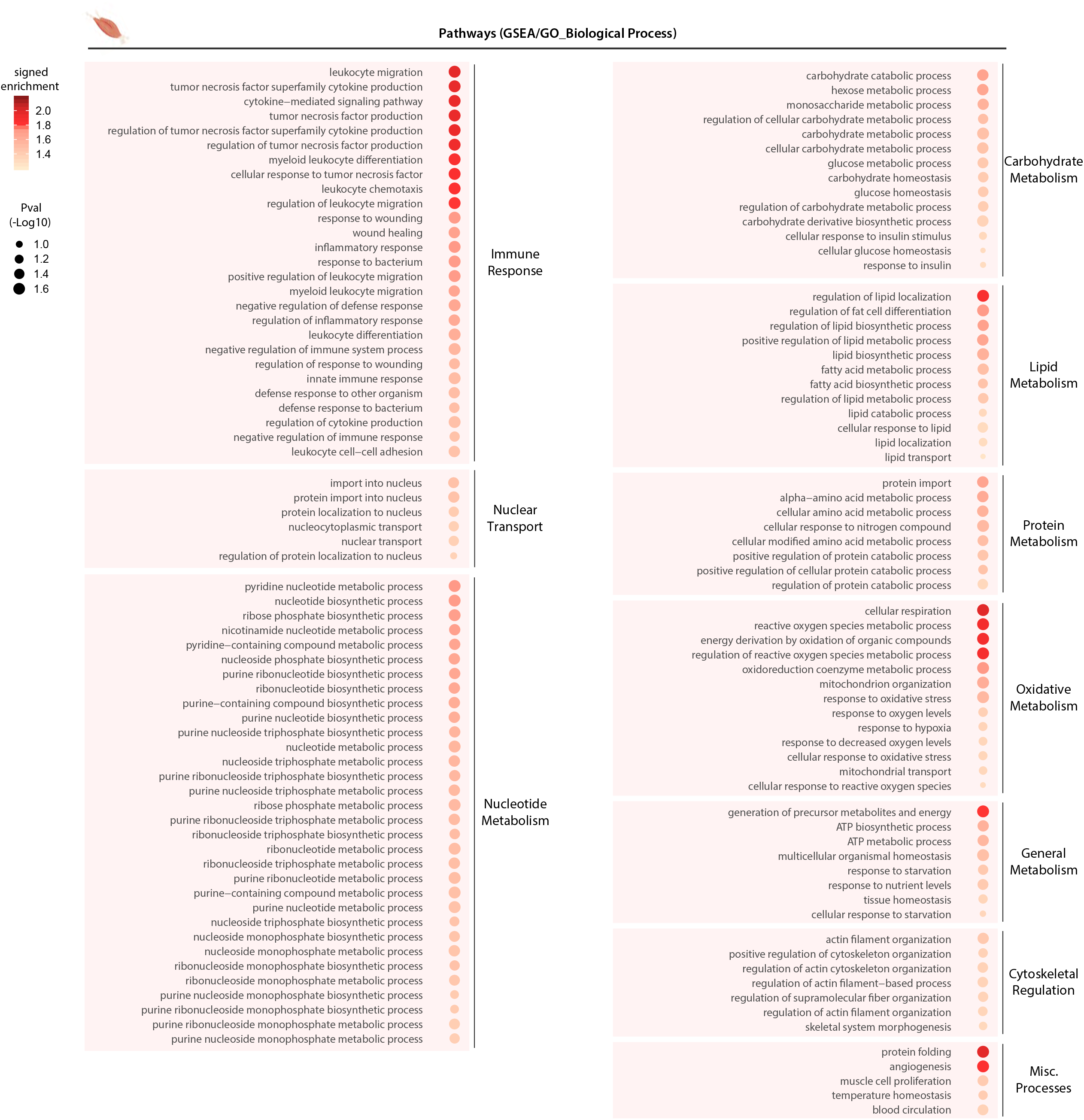
Gene expression analysis on skeletal muscle from exercised MOTS-c-treated old mice. RNA-seq was performed on skeletal muscles from MOTS-c-treated old mice. Balloon plots of biological processes derived from Gene Set Enrichment Analysis (GSEA) using the Gene Ontology (Biological Process) database at a false discovery rate (FDR) < 15% (n=6).

**Fig. S9.**
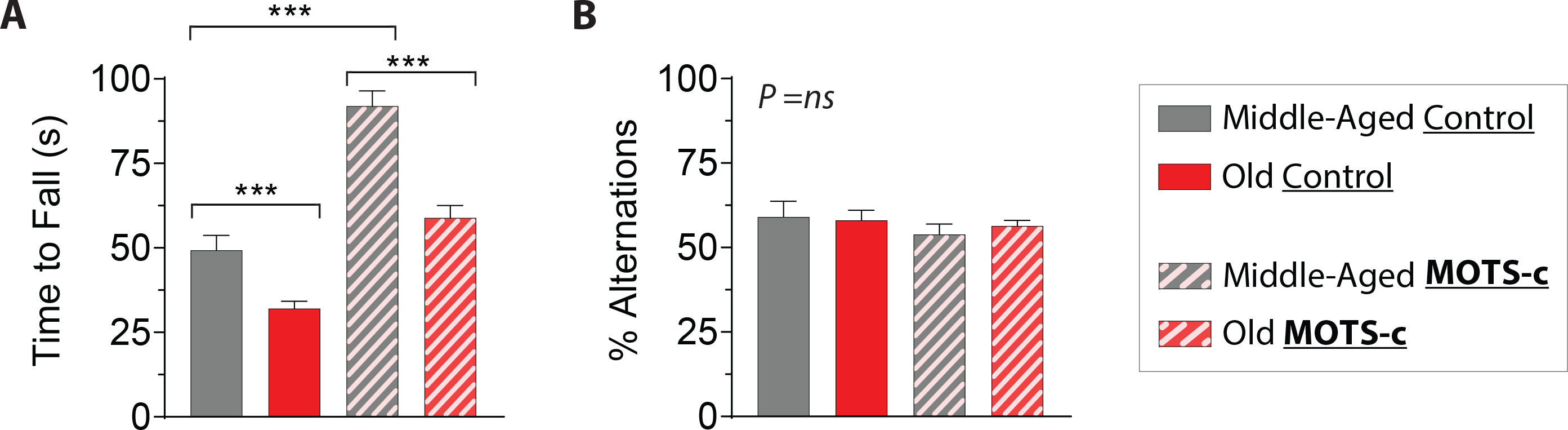
Rotarod and Y-Maze tests in MOTS-c-treated old mice. Middle-aged (14 mo.; n=5-6) and old (24 mo.; n=17-19) mice were treated daily with MOTS-c (15 mg/kg/day; IP) and subject to (**A**) a rotarod test and (**B**) Ymaze test. Data expressed as mean +/− SEM. Student’s t-test. ***P<0.001.

**Fig. S10.**
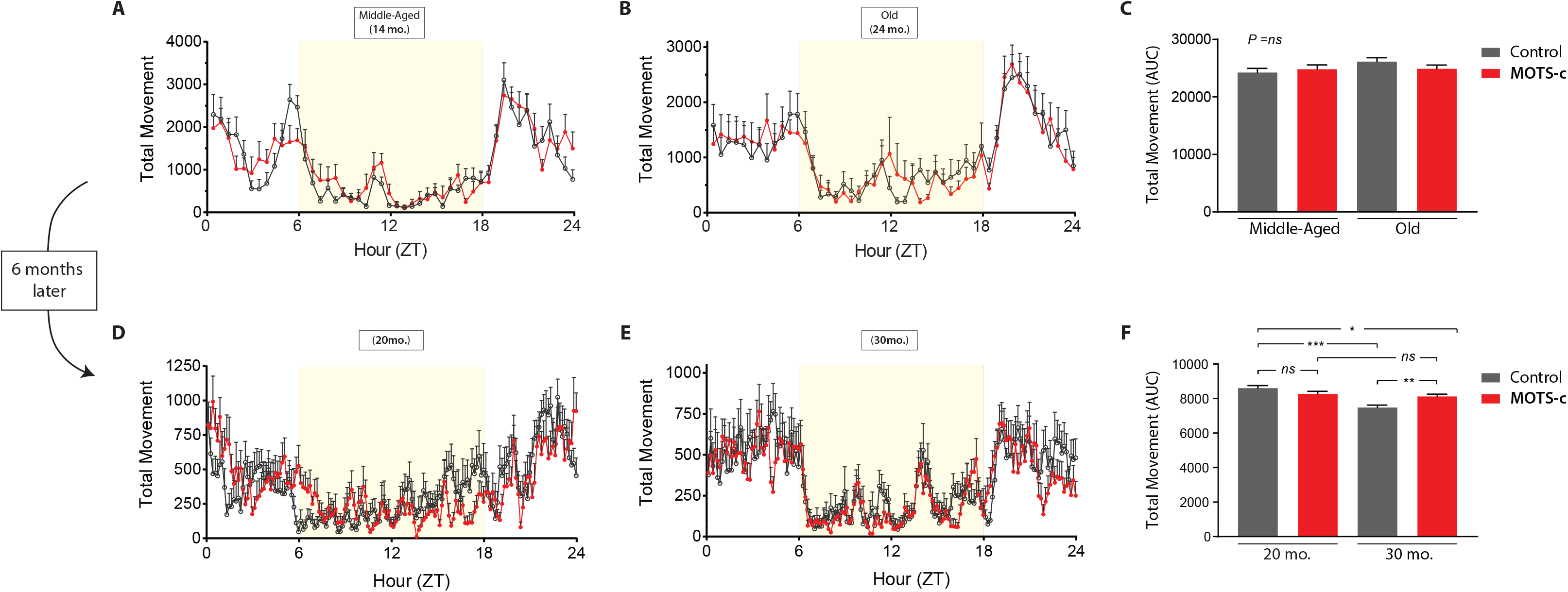
Total physical activity in MOTS-c-treated old mice. Total movement [horizontal and vertical movement (XYZ-axis)] of MOTS-c-treated (**A**) middle-aged (14 mo.) and (**B**) old (24 mo.) mice were continuously measured using metabolic cages throughout the day for three days (n=4). (**C**) The sum of all measured movements is shown. (**D-F**) The procedure was repeated on the same mice after 6 months of LLII MOTS-c treatment. Data expressed as mean +/− SEM of three 24-hour acquisition cycles. Student’s t-test. *P<0.05, **P<0.01, ***P<0.001.

**Fig. S11.**
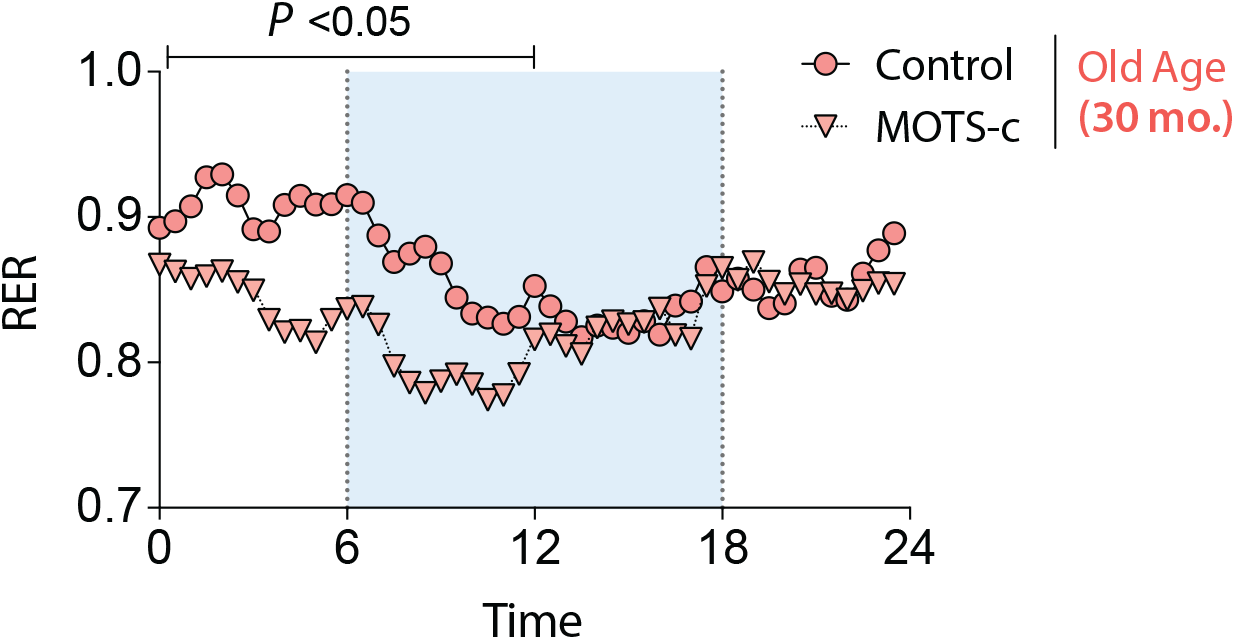
MOTS-c-dependent circadian fuel selection old mice. Respiratory Exchange Ratio (RER) measurements in LLII MOTS-c-treated, or control, old mice (30 mo.; n=4). Shaded region represents daytime (light cycle). Data expressed as mean +/− SEM of three 24-hour acquisition cycles. Two-way ANOVA (repeated measures).

**Fig. S12.**
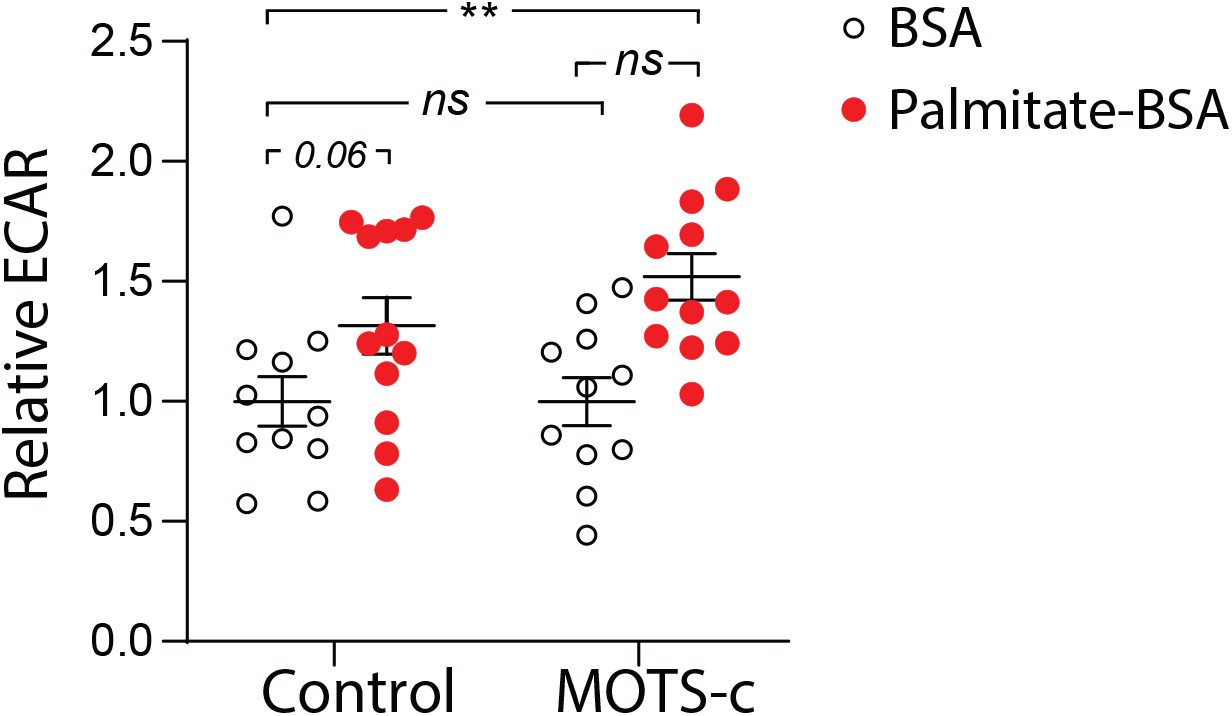
MOTS-c-dependent glycolytic rate in lipid-stimulated mouse myoblasts. C2C12 mouse myoblasts were treated with MOTS-c (10μM) or saline control in nutrient- limited media (n=11-12). Real-time glycolytic flux determined by the extracellular acidification rate was measured using the XF96 Seahorse bioanalyzer. Prior to the start of the assay, nutrient-deprived cells were given either BSA alone or palmitate bound to BSA (palmitate-BSA) to determine the capacity to metabolize fatty acids. Data expressed as mean +/− SEM. Student’s t-test. *P<0.05, **P<0.01, ***P<0.001.

**Fig. S13.**
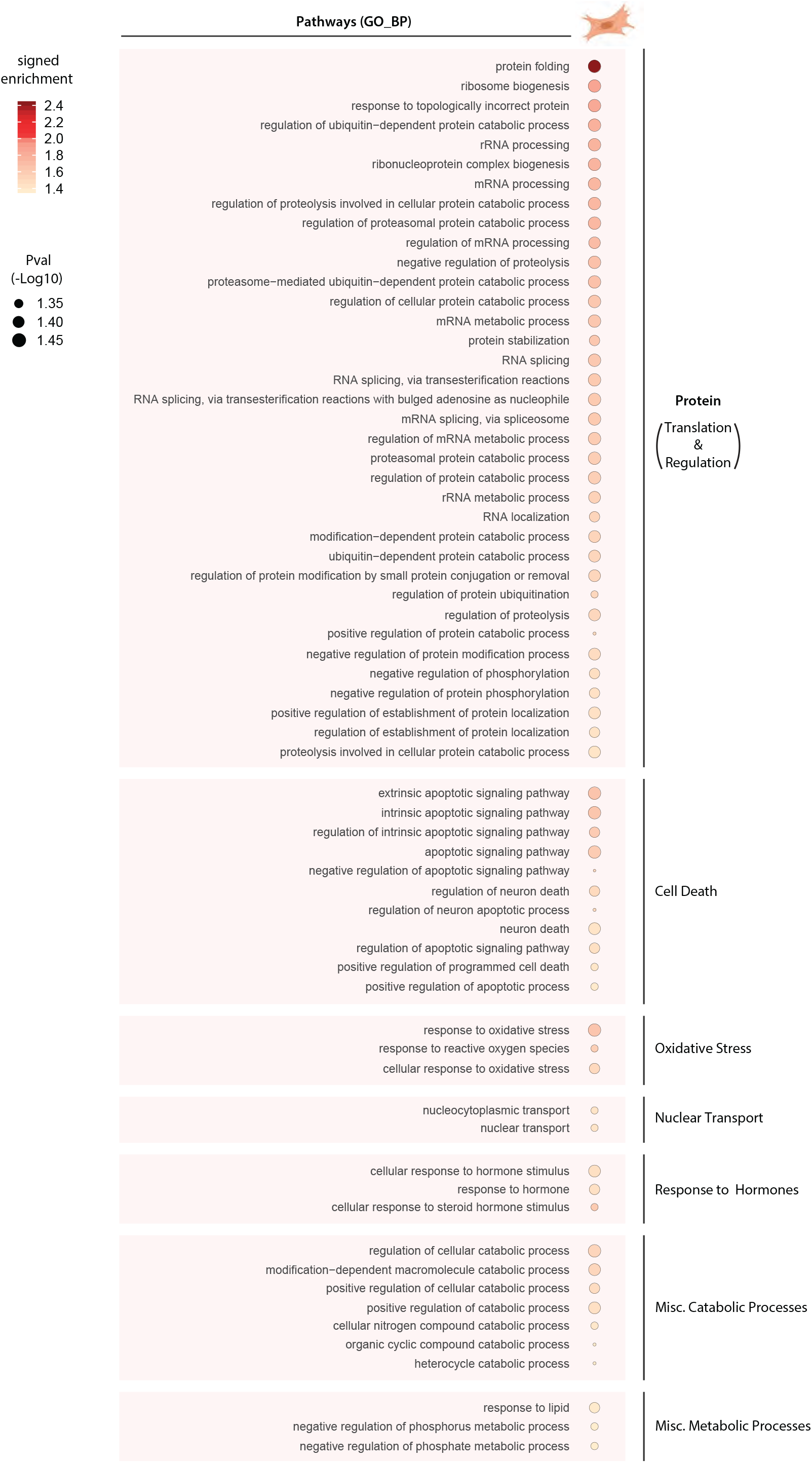
Gene expression analysis on MOTS-c-treated mouse myoblasts under metabolic stress. RNA-seq was performed on C2C12 myoblasts following 48 hours of GR/SD with MOTS-c (10μM) treatment only once initially (n=6). Balloon plots of biological processes derived from Gene Set Enrichment Analysis (GSEA) using the Gene Ontology (Biological Process) database at a false discovery rate (FDR) < 15% (n=6).

